# PROPEL: a high-throughput shear-stress platform reveals organotypic thresholds in endothelial mechano-adaptation

**DOI:** 10.64898/2026.07.22.740104

**Authors:** Terry Ching, Jessica L. Teo, Winita Wangsrikhun, Xiaolei Song, Christopher S. Chen

## Abstract

Endothelial cells exhibit organotypic specialization, yet the tools to decode whether hemodynamic shear shapes this diversity across vascular beds have remained limited. Here we present PROPEL, a tubing-free, magnetic stirrer-driven platform that delivers programmable laminar shear stress (2 to 60 dyn/cm²) to multiple cell types in parallel within the confines of a standard Petri dish, with the modular flexibility to easily introduce different substrate geometries including 3D vessel formats. Using this platform, we profiled six human endothelial subtypes across static, low, intermediate, and high shear. Phenotypic profiling revealed subtype-specific thresholds for alignment, elongation, and Golgi-nuclear polarization, including a dissociation between elongation and polarization in dermal microvascular and saphenous venous endothelial cells at low shear. Bulk RNA sequencing showed conserved transcriptional programs that shift progressively with shear magnitude, including induction of mechanotransduction pathways alongside suppression of proliferative programs. Phenotype-informed transcriptional comparisons further linked alignment transitions to engagement of cytoskeletal and metabolic signatures. In addition to conserved responses, organotypic-specific responses were also observed. Together, these findings establish PROPEL as a scalable platform for multi-endothelial, multi-shear transcriptomic and phenotypic profiling, and reveal that endothelial cells engage shear adaptation along two organotypic axes—a threshold axis governing when a subtype responds and a signature axis governing which the shear stress response—with implications for vascular bed-specific disease susceptibility.

## Introduction

Endothelial cells display distinct molecular identities depending on their vascular bed of origin, with organotypic differences in structure, gene expression, and function^1–3^. Hemodynamic shear stress, a major regulator of vascular health and disease, exhibits some local differences that contribute to focal disease, but cannot explain all observed patterns of disease^4,5^, suggesting the possibility of an interaction between local hemodynamic cues and organotypic endothelial identity contributing to site-specific vulnerability. Endothelial cells depending on their location can experience shear stress spanning a wide physiological range, from approximately 1-6 dyn/cm² in veins to 10-70 dyn/cm² in arteries^6–8^, and distinct endothelial populations are thought to maintain characteristic shear stress “setpoints” to which they are tuned^8,9^. However, whether organotypic endothelial cells interpret shear magnitudes similarly or divergently, and the thresholds at which their phenotypic and transcriptional responses shift, has not been widely investigated.

Addressing these questions requires comparing multiple endothelial subtypes across multiple shear magnitudes under matched conditions, yet existing platforms are not equipped for such multi-parameter studies at sufficiently high throughput to make substantial progress. For instance, syringe- and peristaltic-pump systems deliver precise shear but required a dedicated pump line to each condition and cannot recirculate without added tubing that rapidly becomes untenable^10–13^. Rocking and gravity-driven systems reach plate-scale throughput but impose oscillatory, bidirectional flow rather than defined unidirectional shear^14,15^. Integrated cartridge systems embed perfusion but rely on costly, proprietary consumables that limit scalability^16–19^. Each of these systems requires substantial investment of time, expertise, and resources to adopt, in contrast to, for example, the Petri dish, which has enjoyed widespread adoption. No existing platform combines defined unidirectional shear, high throughput, and low production and adoption cost in a single format, and, more importantly, the ability to access the full physiological range (2– 60 dyn/cm²) across multiple cell types at once. As a result, despite extensive studies on endothelial identity^1,4,20^ and shear-stress mechanotransduction^8,9,21,22^, a systematic, matched comparison across both axes has remained largely out of reach.

To overcome this, we developed Programmable Rotary On-chip Perfusion through an Enclosed Loop (PROPEL), a tubing-free, multichannel platform that integrates into a standard Petri dish. A custom magnetic impeller, actuated by a standard commercial magnetic stirrer, drives continuous, unidirectional, recirculating flow across multiple channels without pumps, tubing, or connectors. Shear magnitude is defined by impeller rotational speed across a 2-60 dyn/cm² range. The simple, low-cost fabrication and scalable channel layout allow testing of multiple cell types and shear conditions in a single, low-cost experiment. Its self-contained format also sustains stable, sterile flow over weeks and yields sufficient biomass for transcriptomic profiling without sample pooling.

To demonstrate these capabilities, we employed PROPEL to investigate whether endothelial cells from different vascular beds might exhibit differential responses to shear stress. We exposed endothelial cells from six vascular beds (brain, skin, aorta, coronary artery, saphenous vein, and umbilical vein) to static, low (2 dyn/cm^2^), intermediate (15 dyn/cm^2^), and high (40 dyn/cm^2^) shear. Next, we profiled phenotypic and transcriptional readouts that are established markers of shear adaptation, including cell alignment, elongation, golgi-nuclear polarization, and induction of the flow-responsive transcription factors KLF2 and KLF4^23–26^. We found that shear thresholds for cellular alignment and polarization differ across vascular beds and changes in alignment were linked to coordinated shifts in biological signatures. Furthermore, increasing shear magnitude led to progressive transcriptional remodeling, suppression of proliferative and induction of mechanotransduction and membrane remodeling pathways. Collectively, these findings establish PROPEL as an accessible platform for organotypic mechanobiology and, using it, show that endothelial cells engage shear adaptation at vascular-bed-specific thresholds and through a graded transcriptional program, with this intitial multi-endothelial, multi-shear dataset made openly available to the community.

## Results

### A tubing-free magnetic impeller drives programmable, recirculating wall shear stress in channels within the confines of a standard Petri dish

PROPEL is a tubing-free, self-contained perfusion platform that generates continuous recirculating flow within a standard 100 mm petri dish, without external pumps, tubing, or connectors. The platform comprises three modular components, a radial fluidic channel array, a magnetic impeller, and a top bracket, that assemble within the dish (Supplementary Fig. 1). The impeller carries an embedded permanent magnet and is actuated by an external magnetic stirrer, allowing flow to be set by rotational speed (RPM) (Fig. 1a; see Methods for fabrication and assembly details). As the impeller rotates, it displaces fluid radially outward, generating a local negative pressure at the channel outlets positioned directly beneath it near the center of the dish (Fig. 1b). This pressure gradient draws fluid inward through the radially arranged fluidic channels, establishing continuous recirculating flow (Fig. 1c). A defined slit (500 µm) between the top bracket and the channel array surface permits outward displacement while suppressing backflow, supplying recirculating fluid to the channel inlets at the outer edge of the dish. The impeller, array, and top bracket are modular and self-contained within the dish (Fig. 1c,d). Cells seeded within the channels thereby experience physiologically relevant wall shear stress under continuous perfusion (Fig. 1d, Supplementary Video 1). Because the radially oriented inlet-outlet design dictates flow, the channels can be easily designed with different geometries and can be designed to integrate 3D cylindrical channels passing through an extracellular matrix gel, supporting engineered 3D vessels (Supplementary Fig. 2).

**Fig. 1.**
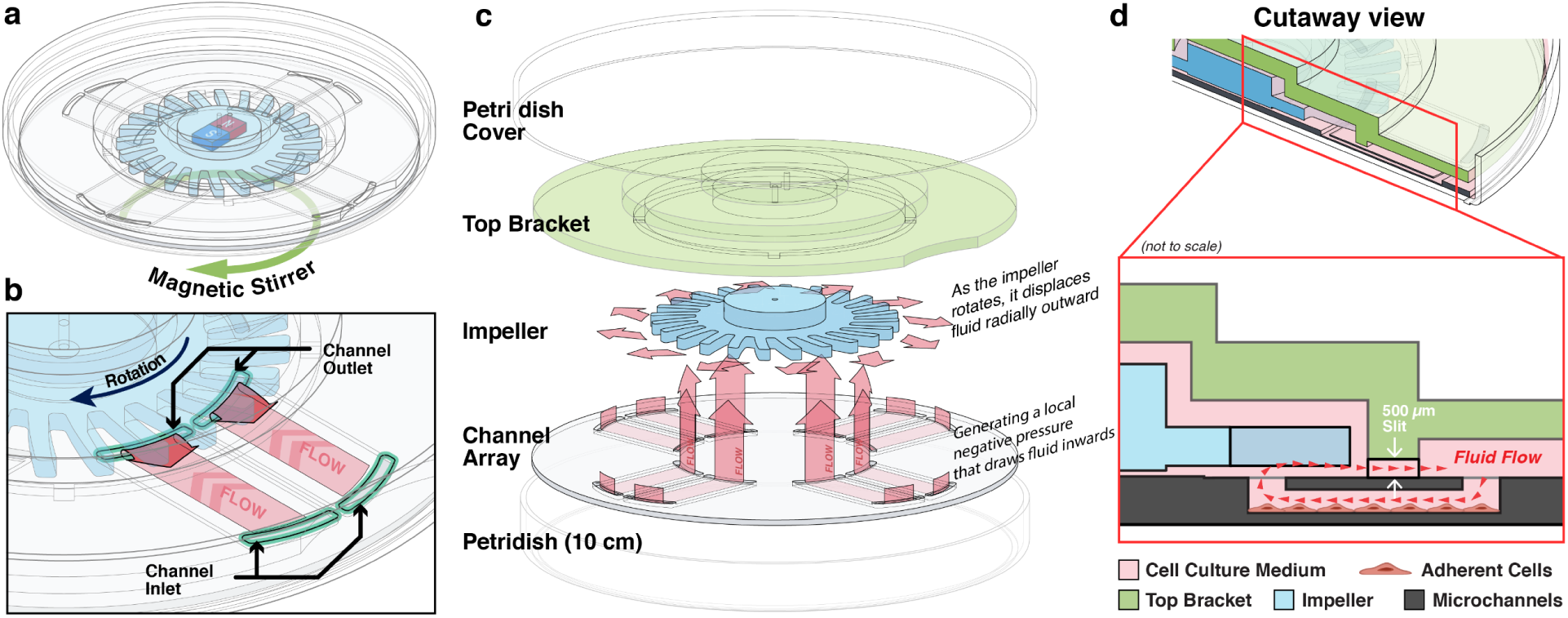
A tubing-free, high-throughput platform for programmable shear stress. (a) Illustration of the impeller, which contains embedded permanent magnets and is magnetically coupled to a commercial magnetic stirrer; impeller rotation, and thus flow rate, is set by the stirrer speed. (b) Schematic of fluid recirculation through the radial channel array: rotation of the impeller displaces fluid outward, drawing it inward through the microfluidic channels to establish continuous recirculating flow (red arrows). (c) Exploded view of the assembled components: top bracket, impeller, channel array, and petri dish. (d) Cutaway view showing endothelial cells seeded within a channel and the direction of fluid flow over the adherent monolayer (red arrow). In all panels, red arrows indicate the direction of fluid flow.

### Wall shear stress is uniform across channels and tunable across 2–60 dyn/cm² by impeller speed and channel dimension

Using particle streak velocimetry (Methods, Supplementary Video 2), we characterized wall shear stress (WSS) across the eight radially arranged channels as a function of impeller speed (Fig. 2a). WSS increased with impeller speed, from 7.3 ± 1.7 to 60.5 ± 4.7 dyn/cm² (200– 700 RPM), with a mean CV of 8.4% across the 300–700 RPM working range (Fig. 2b). Lateral velocity profiles were flat across the channel core, with the no-slip side-wall influence confined to within ∼0.6 mm of each wall (approximately 1.6X of channel height). Hence, majority of the channel experiences uniform, parallel-plate flow (Fig. 2c). Together, these data show that the platform delivers stable, finely tunable shear stress set simply by impeller speed.

**Fig. 2.**
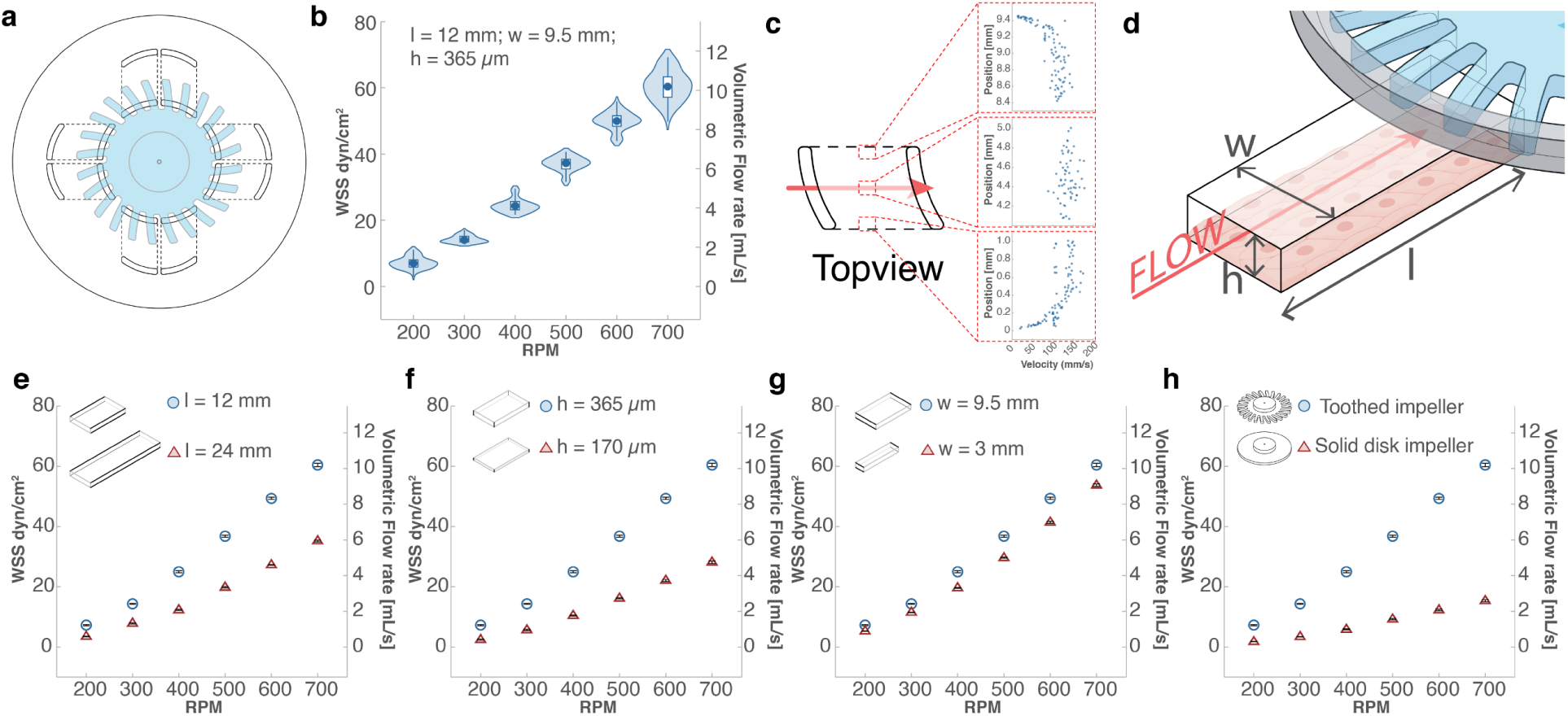
Validation of shear magnitude and spatial uniformity across the platform. **(a)** Top view of the 8-channel array used for flow characterization (L = 12 mm, w = 9.5 mm, h = 365 µm per channel). **(b)** Violin plot of wall shear stress (WSS) across all eight channels, showing the spread between channels; the right axis indicates the corresponding cumulative volumetric flow rate. **(c)** Lateral velocity profiles measured at three positions across a channel (near each side wall and at the center), demonstrating a uniform velocity core flanked by thin near-wall regions. **(d)** Schematic of the channel dimensions (L, w, h) that govern WSS. **(e-h)** Effect of individual geometric and design parameters on WSS, with all other parameters held constant: **(e)** channel length (L = 12 vs. 24 mm), **(f)** channel height (h = 365 vs. 170 µm), **(g)** channel width (w = 9.5 vs. 3 mm), and **(h)** impeller geometry (toothed rotor vs. solid disk). Data in **(e– h)** are shown as mean ± SEM.

We next asked how channel dimensions tune the operating shear (Fig. 2d). Doubling channel length (12→24 mm) approximately halved WSS and reducing channel height (365→170 µm) likewise reduced WSS by about 50% (Fig. 2e,f), both consistent with parallel-plate scaling under a fixed impeller-driven pressure drop^27^. In contrast, narrowing the width (9.5→3 mm), which drops the aspect ratio from ∼26 to ∼8, changed WSS by <10% (Fig. 2g). Finally, replacing the toothed impeller with a smooth cylindrical disk substantially lowered WSS (Fig. 2h), offering an additional, geometry-independent means of accessing lower, more finely controlled shear when required.

### HUVECs recapitulate canonical shear-induced alignment, show KLF2/KLF4 induction, and remain viable under weeks of sustained flow

Having shown that the platform reproducibly delivered uniform and defined shear regimes, we proceeded to test its biological fidelity. To do so, we exposed HUVECs to low (2 dyn/cm^2^) and intermediate (15 dyn/cm^2^) shear stress for 24hrs and assessed for canonical shear-induced changes (Fig. 3a). First, we quantified the expression levels of *KLF2* and *KLF4*, flow sensitive transcription factors involved in endothelial homeostasis^24,28^, using quantitative PCR. Cells cultured under 15 dyn/cm^2^ showed robust upregulation of both transcription factors, whereas cells exposed to 2 dyn/cm^2^ maintained expression levels comparable to those under static conditions (Fig. 3b,c). Next, we examined for shear-induced cell reorientation^26,29,30^ by immunolabeling VE-cadherin, a fiducial marker for cell boundaries, in HUVECs. Cells subjected to 15 dyn/cm^2^ shear showed increased cell alignment (Fig. 3d,e) and elongation (Fig. 3f) along the flow axis (Supplementary Video 3), while cells under 2 dyn/cm^2^ remained morphologically indistinguishable from static conditions. These observations are consistent with prior studies demonstrating that endothelial cells display threshold-dependent shear sensing, where intermediate shear elicits morphological realignment and transcriptional reprogramming^31,32^. Finally, to test whether the platform supports long-term endothelial culture, cells were subjected to 15 dyn/cm^2^ for up to 28 days. Prolonged exposure to shear stress induced alignment in the direction of flow with sustained viability confirmed by Calcein AM staining (Fig. 3g).

**Fig. 3.**
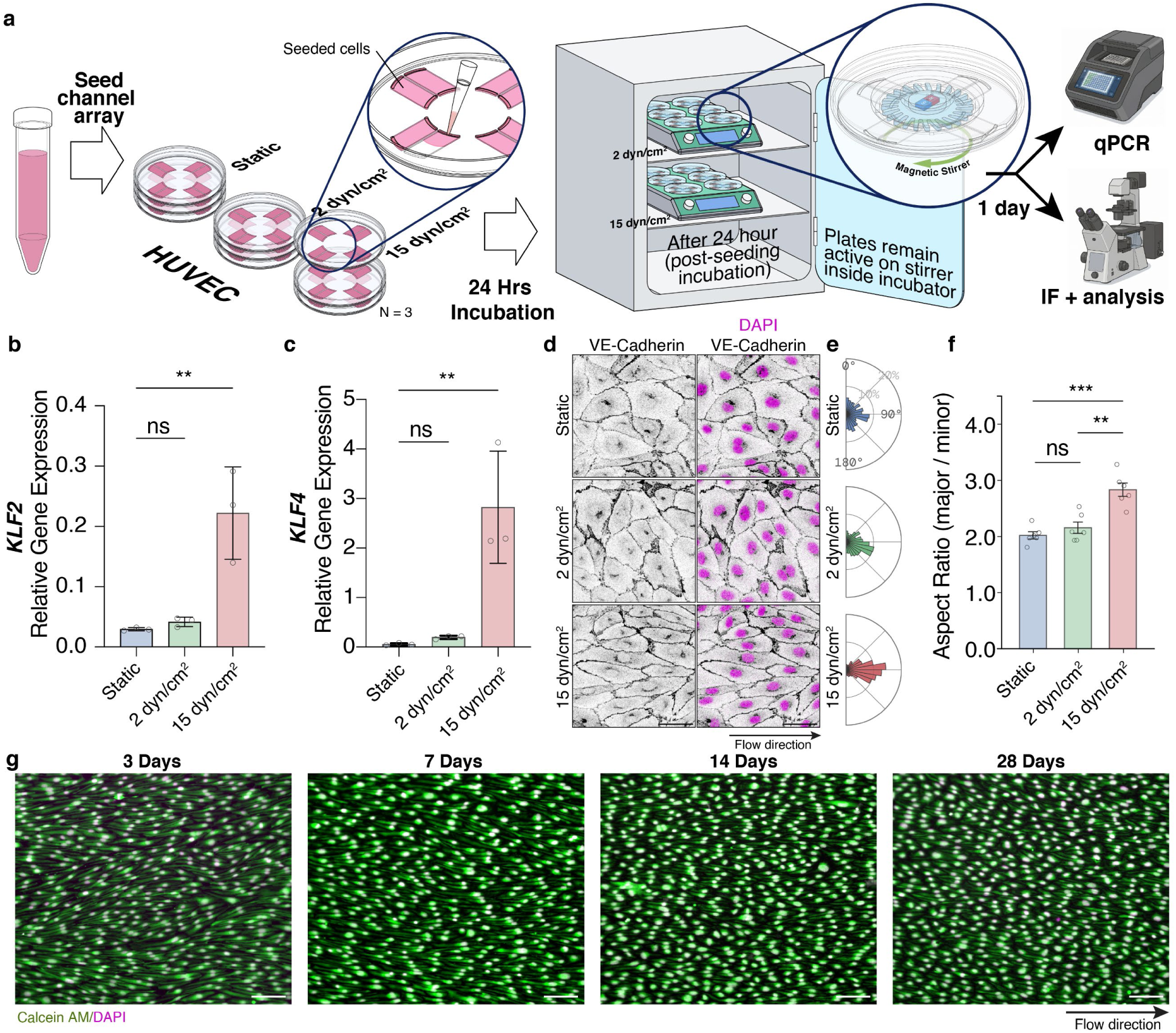
Benchmarking of PROPEL platform with HUVECs. **(a)** Schematic showing experimental workflow. HUVECs were seeded into the channel arrays and maintained in the incubator for 24hrs before subjecting cells to static, 2 dyn/cm², or 15 dyn/cm² conditions. Cells were isolated for qPCR or immunofluorescence analysis 24hrs post-flow. **(b,c)** Relative gene expression of KLF2 **(b)** and KLF4 **(c)** in cells exposed to static, 2 dyn/cm², or 15 dyn/cm² conditions. **(d)** Representative immunofluorescence images of VE-cadherin (left) and VE-cadherin/DAPI overlay (right) of HUVECs exposed to static, 2 dyn/cm², and 15 dyn/cm² conditions. Scale bar = 50 µm. **(e)** Polar histograms of cell orientation angle relative to flow direction. **(f)** Quantification of aspect ratio (major axis/minor axis) of cells subjected to static, 2 dyn/cm², and 15 dyn/cm² conditions. **(g)** Representative Calcein-AM (green) and DAPI (magenta) images of HUVECs subjected to 15 dyn/cm² shear for 3, 7, 14, and 28 days. Scale bar = 100 µm. All data are presented as mean ± s.d. unless otherwise noted; calculated from N=3 independent experiments analysed with one-way ANOVA, Dunnett’s multiple comparisons test. n.s, not significant; *P<0.05, **P<0.01, ***P<0.001, ****P<0.0001.

### Organotypic endothelial subtypes align and polarize at vascular-bed-specific shear thresholds

To demonstrate the feasibility of using this platform for high-throughput investigation of endothelial cell responses to shear stress, we cultured human endothelial cells from six sources: brain (BMVEC), skin (DMVEC), aorta (HAEC), coronary artery (HCAEC), saphenous vein (HsaVEC), and umbilical vein (HUVEC) (Fig. 4a). These cells were exposed to low (2 dyn/cm^2^), intermediate (15 dyn/cm^2^), or high (40 dyn/cm^2^) shear stress for three days before assessing for tissue-specific shear sensitivities. To begin, we examined changes in *KLF2* and *KLF4* gene expression and observed consistent upregulation of both genes across all endothelial subtypes subjected to intermediate and high shear (Fig. 4b,c and Supplementary Fig. 3a). Strikingly, while intermediate and high shear upregulated *KLF2* gene expression to a similar extent across cell subtypes (Supplementary Fig. 3b,d), the induction of *KLF4* was markedly attenuated in DMVECs (Supplementary Fig. 3c,e) compared to other endothelial populations. This is likely due to the higher *KLF4* expression in DMVEC cultured under static condition (Fig. 4c). Additionally, KLF2 consistently showed a lower average induction than KLF4 across different subtypes (Fig. 4b,c). This differential induction illustrates the platform’s sensitivity to subtype-specific transcriptional responses at matched shear magnitude.

**Fig. 4.**
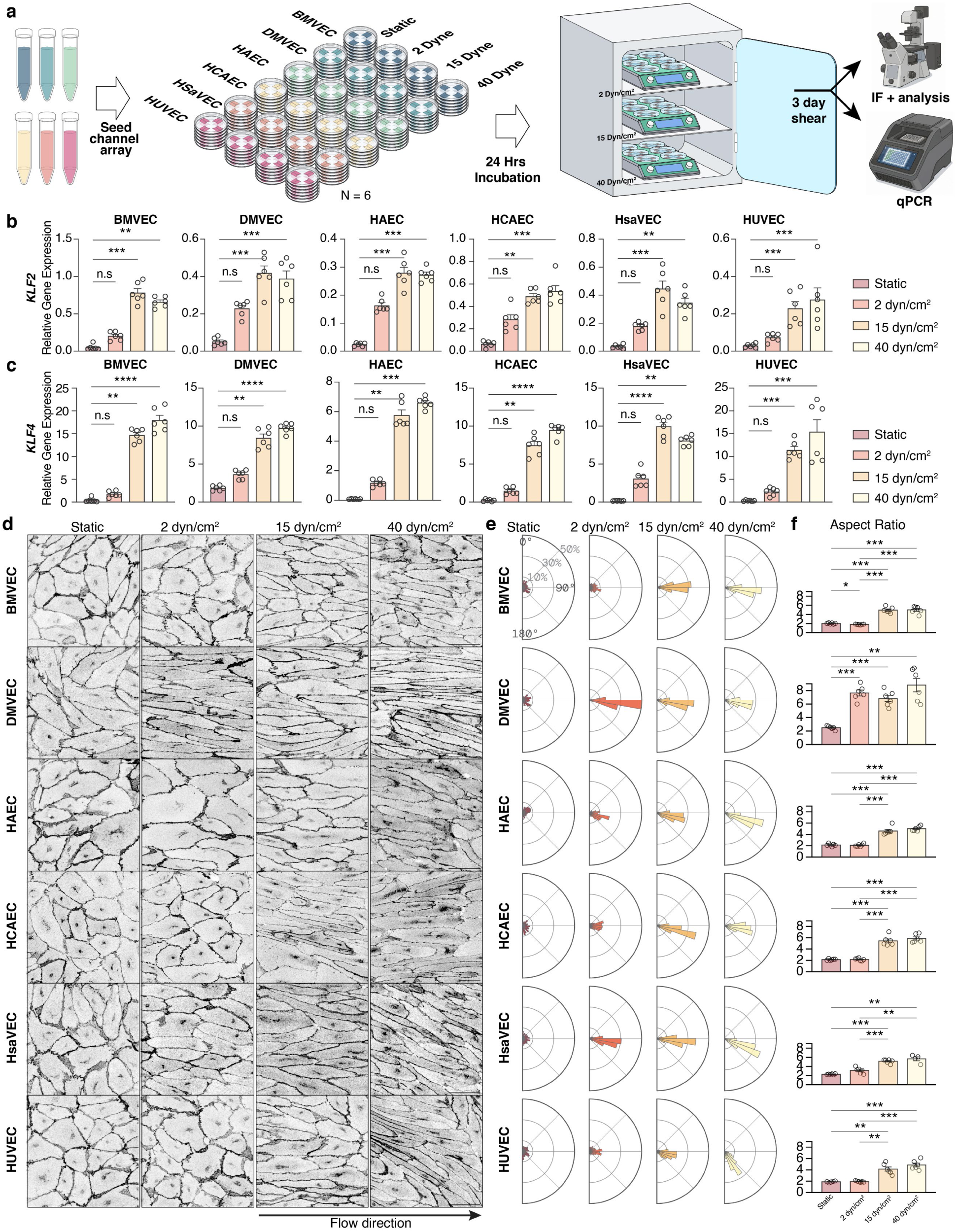
PROPEL reveals subtype-specific phenotypic responses to varying shear magnitudes. **(a)** Schematic showing experimental overview. Six endothelial subtypes (BMVEC, DMVEC, HAEC, HCAEC, HsaVEC, HUVEC) were exposed to static, 2 dyn/cm², 15 dyn/cm², and 40 dyn/cm² conditions for 3 days. Cells were then isolated for qPCR or immunofluorescence. **(b,c)** Relative gene expression of KLF2 **(b)** and KLF4 **(c)** in six endothelial subtypes subjected to static, 2 dyn/cm², 15 dyn/cm², and 40 dyn/cm² conditions for 3 days (N=6). **(d)** Representative immunofluorescence images of VE-cadherin in each of the six endothelial subtypes under static, 2 dyn/cm², 15 dyn/cm², and 40 dyn/cm² conditions for 3 days. **(e)** Polar histograms of cell orientation angle relative to flow direction for each subtype under static, 2 dyn/cm², 15 dyn/cm², and 40 dyn/cm² conditions for 3 days. **(f)** Quantification of cell aspect ratio (major axis/minor axis) for each subtype across the four shear conditions. All data are presented as mean ± s.e.m. unless otherwise noted; calculated from N=6 independent experiments analyzed with one-way ANOVA, Dunn’s multiple comparisons test. n.s, not significant; *P<0.05, **P<0.01, ***P<0.001, ****P<0.0001. Scale bar = 50mm.

Beyond *KLF2* and *KLF4*, we profiled a panel of shear-responsive genes known to be either induced or repressed by laminar shear^25,33–35^. Consistent with canonical laminar shear response, *NOS3* and *THBD* were induced at intermediate and high shear across all six subtypes (Supplementary Fig. S3a). Conversely, the proinflammatory and vasoactive genes *CCL2*, *VCAM1*, *SELE*, and *ET-1* were repressed at intermediate and high shear relative to static and low-shear conditions (Supplementary Fig. S3a). This bidirectional response demonstrates that the platform captures both arms of the canonical shear-adaptive transcriptional program.

When comparing shear-induced cell reorientation across subtypes, DMVECs and HsaVECs displayed enhanced cell alignment and elongation along the flow axis at the low shear stress of 2 dyn/cm^2^ (Fig. 4D, E, and F). In contrast, BMVECs, HAECs, HCAECs, and HUVECs remained morphologically similar to static controls (Fig. 4D, E, and F). Consistent with canonical flow responses, intermediate and high shear promoted cell alignment and elongation across all subtypes (Fig. 4d,e,f). Notably, at 40 dyn/cm^2^, HUVECs showed a near perpendicular orientation relative to the flow axis. This is consistent with reports showing that HUVEC alignment is restricted to a 10-20 dyn/cm^2^ set point and reverses outside this range^9,36^. In addition to alignment and elongation, intermediate shear has been reported to induce endothelial polarization against the direction of flow, with the golgi apparatus localizing upstream of the nucleus^22^. However, the direction of polarization has been reported to vary across vascular beds and experimental contexts^22,37^. Hence, we examined golgi-nuclear polarization, to determine whether it tracked with alignment across endothelial subtypes. While endothelial cells cultured under static and low shear conditions remained randomly polarized, at intermediate shear stress, endothelial cells displayed golgi-nuclear reorientation in the direction of flow, except for BMVEC which showed golgi positioning both upstream and downstream of the nucleus (Supplementary Fig. 4a,b). At high shear stress, all endothelial subtypes were polarized, with their golgi positioned upstream of the nucleus against the direction of flow (Supplementary Fig. 4a,b). Interestingly, only DMVECs showed polarization at 2 dyn/cm^2^ (Supplementary Fig. 4a,b), despite both DMVECs and HsaVECs displaying alignment and elongation along the flow axis at low shear (Fig. 4d,e,f). Alignment and Golgi–nuclear polarization can thus be uncoupled in an organotypic manner: both low-shear responders align, yet only dermal microvascular cells repolarize. Together, these findings demonstrate the platform’s suitability for high throughput phenotypic assessment of shear responses across multiple cell types and magnitudes in parallel, at a scale not readily achievable with lower throughput systems.

### Progressive transcriptional remodeling and biological signature shifts with shear magnitude

In addition to quantitative analysis of endothelial phenotypes, the platform can also support omics profiling under controlled hemodynamic conditions. Therefore, we conducted bulk RNA sequencing on all six endothelial subtypes subjected to static, low, intermediate, and high shear stress to identify common shear-responsive genes, genes unique to specific shear magnitudes, and biological signatures associated with endothelial alignment phenotypes (Fig. 5a). Because these subtypes are cultured in their own subtype-specific medium, shear-responsive patterns observed consistently across all subtypes are likely to reflect genuine biological signal rather than a media- or cell-line specific artifact. To begin, we sought to validate our dataset, by comparing the transcriptional response of HUVECs subjected to intermediate shear stress against two publicly available datasets (GSE151867^38^ and GSE294621^39^). 21-44% of our differentially expressed genes (Supplementary Fig. 5a) were present in the public dataset and spearman correlation of log2 fold-changes across all shared genes between datasets showed r = 0.225 and r = 0.265 respectively (Supplementary Fig. 5b,c). While concordance across all shared genes was modest, consistent with differences in platform, shear magnitude, and sequencing depth, differentially expressed genes in both datasets revealed strong concordance (GSE151867: r = 0.798, n = 205; GSE294621: r = 0.518, n = 269) (Supplementary Fig. 5d,e), indicating that high-confidence shear-responsive genes are highly reproducible across datasets. Canonical flow-responsive genes including *KLF2*, *KLF4*, *NOS3*, and *THBD* were consistently and significantly upregulated across all three datasets (Supplementary Table 1), confirming that laminar shear stress-induced transcriptional program is detected in our dataset.

**Fig. 5.**
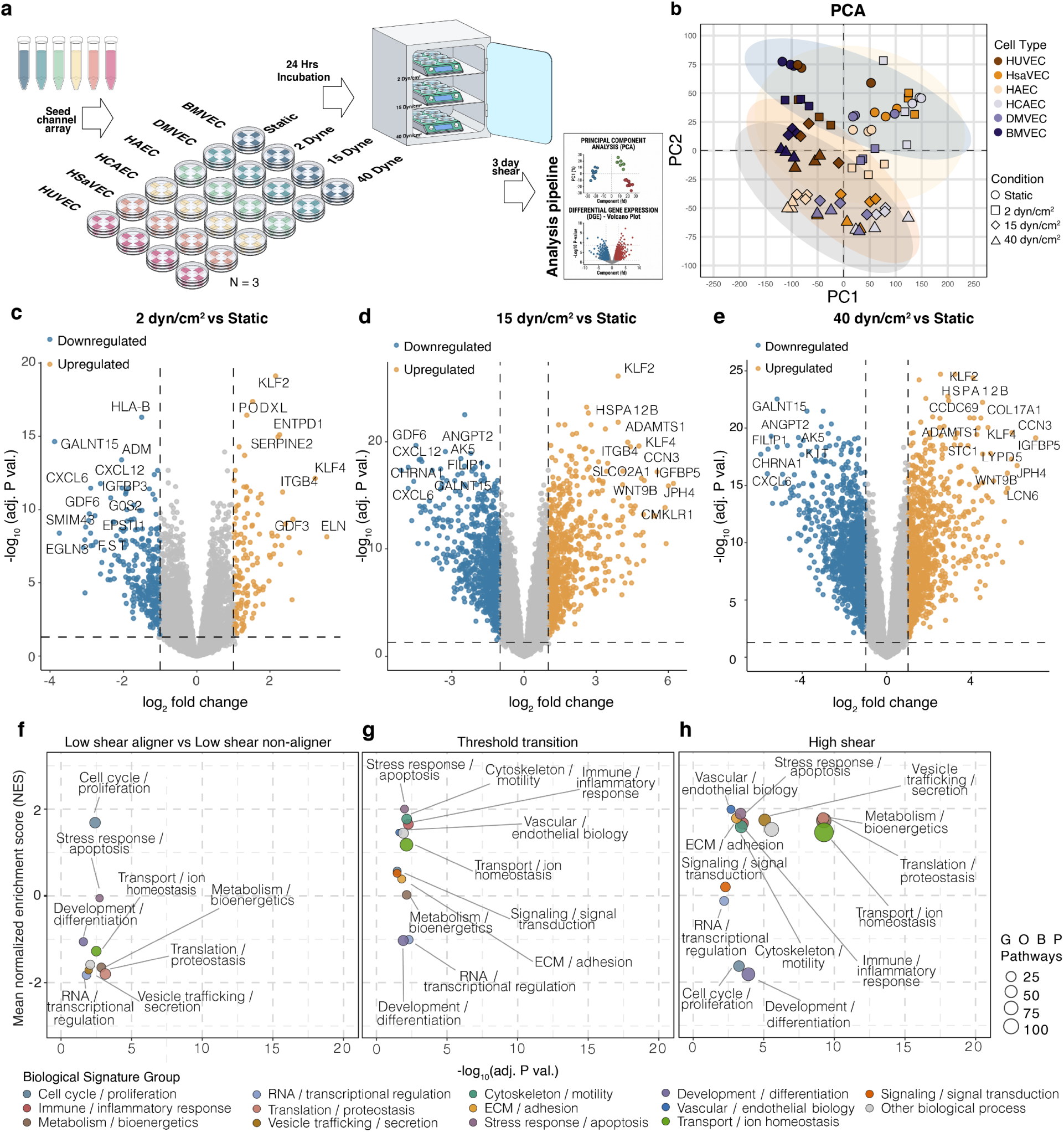
Transcriptomic profiling reveals graded responses to shear stress. **(a)** Schematic of experimental overview for bulk RNA sequencing of six endothelial subtypes (BMVEC, DMVEC, HAEC, HCAEC, HsaVEC, HUVEC) under static, 2 dyn/cm², 15 dyn/cm², and 40 dyn/cm² conditions. **(b)** Principal component analysis of RNA-seq samples, colored by endothelial subtype and shaped by shear condition. Static (blue), 2 dyn/cm² (yellow), 15 dyn/cm² (orange), and 40 dyn/cm² (grey) ellipse represent the 95% confidence interval. **(c-e)** Volcano plots of differentially expressed genes for 2 dyn/cm² vs. static **(c)**, 15 dyn/cm² vs. static **(d)**, and 40 dyn/cm² vs. static **(e)** comparisons. Upregulated genes (orange) and downregulated genes (blue) are defined by adjusted P value and log_2_ fold change thresholds (dashed lines); Top 20 genes are labeled. **(f-h)** Gene Set Enrichment Analysis of Gene Ontology Biological Process pathways, grouped into biological signature categories for low-shear aligner vs. low-shear non-aligner comparison **(f)**, threshold transition **(g)**, and high-shear condition **(h)**. Bubble size indicates the number of GOBP pathways contributing to each signature group.

Next, principal component analysis (PCA) of global transcriptomic profiles of our dataset showed that endothelial cells segregated based on both vascular bed identity and shear exposure. Interestingly, we observed progressive shifts along PC2 across low-, intermediate-, and high-shear conditions, indicating stepwise mechanoadaptation transition (Fig. 5b). To identify shear-responsive genes at low, intermediate, and high shear relative to static culture, we performed pairwise differential gene expression analyses. Several genes associated with immune response (*CXCL6*^40^*, CXCL12*^41^*, HLA-B*^42^) and vascular dysfunction (*ANGPT2*^43,44^*, EPSTI1*^44^) were significantly downregulated in response to shear stress. Conserved endothelial shear-responsive genes, such as *KLF2* and *KLF4*^24,28^, were consistently upregulated, regardless of shear magnitude, whereas *CXCL6* was significantly downregulated across these conditions (Fig. 5c,d,e). Exposure to low shear reduced the expression of genes associated with vasodilation (*ADM*^45,46^*, FST*^47^) and genes associated with pro-survival signaling (*G0S2*^48^*, IGFBP3*^49^) (Fig. 5c). Notably, several genes associated with matrix modeling *(ADAMTS1*^50,51^*, CCN3*^52,53^) and pro-survival signaling *(HSPA12B*^54^*, IGFBP5*^55^) were consistently upregulated under both intermediate and high shear conditions, whereas *FILIP1*^56,57^, associated with cytoskeletal remodeling, was consistently downregulated, indicating shared transcriptional features between intermediate and high shear states (Fig. 5d,e). At high shear magnitudes, genes associated with stress response *(LCN6*^58^*, STC1*^59–61^) were significantly upregulated (Fig. 5e). Collectively, these findings highlight the presence of both conserved endothelial shear-responsive genes and genes enriched at specific shear magnitudes.

To determine whether these transcriptional changes reflect discrete condition-specific responses or a continuous dose-dependent program, we performed Spearman correlation analysis between normalized gene expression and ranked shear conditions across all four magnitudes. Genes satisfying a Spearman |ρ| ≥ 0.9 threshold and pairwise differential expression significance (FDR < 0.05) were classified as high-confidence shear-magnitude-dependent genes. In total, 1915 shear-induced and 2197 shear-repressed genes met these criteria, with individual gene trajectories showing directional responses across the full shear range and across six endothelial subtypes (Supplementary Fig. 5f). Within this gene set, we further identified progressive genes (Supplementary Fig. 5g,h), whose expression continued to change between 15 and 40 dyn/cm^2^. These graded transcriptional patterns suggest that endothelial response to flow occurs through progressive transcriptional transitions. The complete list of progressive genes is provided in Supplementary Table 2.

To determine whether shear-magnitude-dependent transcriptional changes converged on coordinated biological programs, we conducted Gene Ontology Biological Process (GO BP) gene set enrichment analysis across all shear conditions. Low shear enriched pathways associated with cellular organization and membrane remodeling, suggesting initial flow adaption (Supplementary Fig. 5i). Intermediate and high shear conditions displayed enrichment in pathways related to membrane targeting and organization, endothelial migration, and oxidative stress responses (Supplementary Fig. 5j,k). Crucially, intermediate shear conditions also showed enrichment of pathway associated with response to fluid shear stress, indicating the activation of canonical endothelial mechanotransduction gene sets related to flow (Supplementary Fig. 5j). Across all shear conditions, pathways related to cell cycle were negatively enriched (Supplementary Fig. 5i,j,k).

Next, we examined whether the endothelial alignment phenotypes observed at varying shear magnitudes were associated with distinct shifts in biological signatures. To test this, we performed three phenotype-informed biological contrasts: low-shear aligners versus low-shear non-aligners, threshold transition response at intermediate shear, and high shear exposure. Enriched GOBP pathways were consolidated into broader biological signatures to enable a system-level interpretation of endothelial shear adaptation (Fig. 5f,g,h). Full differential expression and pathway enrichment results for all biological contrasts are provided in Supplementary Tables 3-12. The response of low-shear aligners was characterized by a relative suppression of translation/proteostasis and metabolism/bioenergetics-associated signatures, alongside apparent enrichment of cell cycle/proliferation-associated pathways (Fig. 5f). Stratified analysis within aligners and non-aligners confirmed that the reduced translation/proteostasis and metabolism/bioenergetics signatures are genuine transcriptional remodeling in low-shear-aligning populations, while the apparent proliferative enrichment was due to stronger suppression of cell cycle-related pathways in non-aligners rather than absolute induction in aligners (Supplementary Fig. 5l,m). Endothelial populations that failed to align under low shear stress but did so at intermediate shear stress displayed enriched signatures associated with cytoskeleton/motility, stress response/apoptosis, and immune/inflammatory response (Fig. 5g). In high-shear exposure, pathways related to metabolism/bioenergetics and transport/ion homeostasis was enriched (Fig. 5h).

## Discussion

Using PROPEL to profile six endothelial subtypes across four shear magnitudes, we find that endothelial shear-adaptive responses are not uniformly engaged: subtypes cross into an adaptive state at vascular-bed-specific thresholds, and co-engage different combinations of outputs once a given threshold is reached. The platform that enabled this matched comparison is itself low-cost, scalable, and high-throughput, addressing a gap left by pump-, rocker-, and cartridge-based shear systems and yielding a multi-endothelial, multi-shear phenotypic and transcriptomic dataset. We make this system openly available to the community through open source designs for fabrication and adoption.

Cellular alignment, golgi-nuclear polarization, and KLF2/KLF4 induction are canonical readouts that are routinely measured in endothelial cell cultures exposed to physiological shear to define the flow-conditioned endothelial phenotype^21,62,63^. At a low shear magnitude of 2 dyn/cm^2^, we found that both DMVECs and HsaVECs engage in cellular alignment unlike the remaining four subtypes. Notably, the two low shear-responsive subtypes also differed in the outputs they engaged: DMVECs showed concomitant golgi-nuclear repositioning, while HsaVEC aligned without it. As a single donor was profiled per subtype in this dataset, this divergence could be attributed to a difference in endothelial identity or donor-to-donor variability. Resolving this will require expanding each subtype to a multi-donor panel, a scale-up within the platform’s capabilities.

Together, these observations point to two separable axes of organotypic shear adaptation: a threshold axis that sets the shear magnitude at which a given subtype first responds, and a signature axis that sets which outputs—alignment, elongation, and Golgi– nuclear polarization—it co-engages once that threshold is crossed. This logic extends the prevailing shear-stress ‘setpoint’ model, in which each endothelium is tuned to a characteristic magnitude: rather than a single tuned value, our data suggest that setpoints are graded across vascular beds and that the phenotypic program engaged at a given magnitude is itself organotypically specified. Because vascular pathologies localize to particular beds and even particular regions of a vessel, such organotypic differences in how a shared hemodynamic input is interpreted could contribute to site-specific disease susceptibility.

Next, our data shows that a considerable portion of the transcriptional response to shear is graded rather than discrete. PCA analysis identified progressive shifts in global transcriptomic state across low, intermediate, and high shear while pathway enrichment analysis identified membrane remodeling programs at low shear, canonical mechanotransduction signatures at intermediate shear, and protein stability programs at high shear. These observations indicate that endothelial shear adaptation unfolds through a progressive sequence in which distinct biological pathways are engaged with increasing shear magnitude.

Apart from endothelial mechanobiology, the platform is broadly applicable to any adherent cell type that experiences flow. For instance, intestinal epithelial cells and renal tubular cells each experience physiologically relevant luminal shear that governs barrier integrity and epithelial organization^64,65^. While shear stress is disrupted in disease in both contexts^66,67^, how changes in shear magnitude regulate disease progression remain understudied. In addition, the low cost and high throughput design positions PROPEL as a tool for drug screening under physiologically relevant flow, enabling pharmacological assessment of vasoactive, anti-inflammatory, or barrier-modulating compounds in hemodynamic conditions that more accurately reflect vessel wall environments. In conclusion, we developed a platform that supports prolonged, non-toxic endothelial culture with programmable shear stress reaching up to 60 dyn/cm^2^. By making defined, physiological shear broadly accessible, PROPEL—together with the multi-endothelial, multi-shear dataset it generates—provides an openly available foundation for the vascular biology, mechanobiology, and bioengineering communities. We anticipate that pairing this accessibility with the organotypic, threshold-and-signature logic uncovered here will deepen understanding of how hemodynamic forces shape endothelial responses in health and disease and motivate the next generation of studies into vascular bed-specific disease susceptibility.

## Methods

### Cell Culture

To support cell adhesion and culture in the channels, channel surfaces were functionalized. Arrays were treated with plasma cleaner (PDC-001, Harrick Plasma, NY, USA) for 6 min to activate the surface, then coated with polydopamine (2 mg/mL dopamine hydrochloride (cat. no. H8502, Sigma-Aldrich, MA, USA) in 1× TBS, pH 8.5) for 40 min. Devices were rinsed three times in deionized water, dried, and sterilized under UV in a biosafety hood for 20 min prior to cell seeding. Six primary human endothelial cell types were used in this study: human umbilical vein endothelial cells (HUVECs), human saphenous vein endothelial cells (HsaVECs), human brain microvascular endothelial cells (BMVECs), human dermal microvascular endothelial cells (DMVECs), human coronary artery endothelial cells (HCAECs), and human aortic endothelial cells (HAoECs). HUVECs (Lonza, cat. no. C2519A) and HAECs (Lonza, cat. no. CC-2535) were cultured in Endothelial Cell Growth Medium-2 BulletKit (EGM™-2 BulletKit; Lonza, cat. no. CC-3162), which comprises Endothelial Cell Basal Medium-2 (EBM™-2; Lonza, cat. no. CC-3156) supplemented with the EGM™-2 SingleQuot™ supplement pack (Lonza, cat. no. CC-4176). HsaVECs (PromoCell, cat. no. C-12231) were cultured in Endothelial Cell Growth Medium (PromoCell, cat. no. C-22110). BMVECs (CellSystems, ACBRI 376), DMVECs (Lonza, cat. no. CC-2516), and HCAECs (Lonza, cat. no. CC-2585) were maintained in EGM™-2 MV Microvascular Endothelial Cell Growth Medium-2 BulletKit (Lonza, cat. no. CC-3202), which comprises Endothelial Cell Basal Medium-2 MV (EBM™-2 MV; Lonza, cat. no. CC-3156) supplemented with the EGM™-2 MV SingleQuot™ supplement pack (Lonza, cat. no. CC-4147). Cells were used at passages 2 to 8. All cells were maintained at 37 °C in 5% CO_2_ in a humidified incubator. Cell line authentication (performance testing, differentiation capacity, and STR profiling) was provided by Lonza and PromoCell at the time of purchase.

### Fabrication of top bracket, impeller, and standard channel array

All components were designed to fit within a 100 mm petri dish (Corning) and fabricated from poly(methyl methacrylate) (PMMA) sheets of varying thickness (0.25, 2, and 3 mm). Throughout, 2 mm and 3 mm sheets were cut on a laser cutter (Fusion Edge, Epilog Laser, Golden, CO, USA), 0.25 mm sheets on a cutting plotter (Silhouette Cameo 5, Silhouette America, Utah, USA), and layers were solvent-bonded with chloroform. The top bracket was cut according to the patterns in Supplementary Fig. 1B. To form its spacing features, two 0.25 mm sheets were first bonded into a 0.5 mm layer, which was then cut with an X-Acto knife into ∼2 × 2 mm squares; all layers were assembled as illustrated in Supplementary Fig. 1B. The bracket was fitted with a central shaft that held the impeller in concentric alignment and maintained a defined 500 µm slit between the impeller and the array surface (Fig. 1C), sized to permit outward fluid displacement during rotation while suppressing backflow. The central shaft was made from a 20-gauge blunt needle, truncated and crimped at one end with a wire cutter. The impeller was cut and assembled from 0.25, 2, and 3 mm PMMA layers as illustrated in Supplementary Fig. 1C. A 10 µL droplet of clear resin (Formlabs, MA, USA) was pipetted onto its underside to form a bulge and cured under a UV lamp (Form Cure V1, Formlabs, MA, USA) for 5 min at room temperature. This cured resin bulge serves as a low-friction pivot point on which the impeller spins. 3 mm (cat. no. 8560K239) and 0.25 mm (cat. no. 4076N11) PMMA sheets were purchased from McMaster-Carr (IL, USA). 2 mm PMMA sheets (cast plexiglass), 20-gauge blunt needle were purchased from Amazon (WA, USA). Fluidic channel arrays were fabricated by xurography of polyethylene terephthalate (PET) film and double-coated tape (ref). The flow channel layer was formed by laminating two 170 µm double-coated tape layers (3M 9495LE, 3M, Minnesota, USA) onto both sides of a ∼25 µm PET carrier film (#48-1F-OC, 0.001″, CS Hyde, Illinois, USA), giving a total channel height of 365 µm. This stack was cut on a cutting plotter (Silhouette Cameo 5, Silhouette America, Utah, USA) according to the pattern in Supplementary Fig. 6A separate ∼100 µm PET film (#48-4F-OC, CS Hyde, Illinois, USA) cut on the same plotter, was positioned over the channel layer to form the ceiling, and glass coverslips were applied below to seal the channels (Supplementary Fig. 6A). Assembled devices were bonded overnight in a 100 °C oven.

### Magnetic stirrer modification

Multi-position magnetic stirrer blocks (MMS-6Pro, Joan Lab, Zhejiang, China) were used to drive the impellers. Each block contains six independent, asynchronous, programmable stirrer positions, and depending on incubator size and rack configuration, two to three blocks were placed inside the incubator (Supplementary Fig. 7A). Because the blocks generate heat during prolonged operation, three modifications were made to protect the cultures. First, to improve cooling, slots were cut into the sides of each block with a universal snip, and two to three 60 mm USB brushless fans (UMLIFE computer PC fans) were mounted with double-sided tape over the slots to circulate air across the internal brushless motors and motor drivers (Supplementary Fig. 7B). Second, to reduce radiative heat transfer to the dish, 9 × 9 cm aluminum foil squares were taped onto the block directly above each motor (Supplementary Fig. 7B). Third, silicone strips (Dragon Skin FX-Pro, Smooth-On, PA, USA), cast at ∼4 mm thickness and cut to size, were placed over each stirrer position (Sup. Fig. XC); these provided friction to hold the dish in place and elevated it off the block surface to reduce conductive heat transfer.

### Measurement of flowrate and calculation of WSS

Volumetric flow rate and wall shear stress (WSS) were quantified by particle streak velocimetry. To image the channels while the impeller was spinning, a programmable magnetic stirrer (MS5S, Joan Lab, Zhejiang, China) was dismantled down to its brushless motor and driving electronics (PCB, motor driver, LCD display, and control knob). The motor was mounted on a three-legged MakerBeam frame (Utrecht, Netherlands) whose legs rested directly on the petri dish lid, positioning the motor above the impeller to drive it magnetically from above. The dish was mounted on the microscope stage using an adjustable petri dish holder, allowing the channels to be imaged through the objective below while the impeller was actuated from above (Supplementary Fig. 8A). Channels were perfused with a suspension of 4 µm fluorescent beads (cat. no. T7283, Invitrogen, CA, USA), and images were acquired at a fixed 1 ms exposure such that each bead appears as a streak whose length equals the distance traveled in 1 ms; bead velocity was therefore obtained directly as streak length divided by exposure time. The focal plane was set to the channel mid-height (h/2), where the parabolic flow profile reaches its maximum (centerline) velocity, *v*_*max*_, and only beads tracked near the lateral center of the channel were used, away from the side walls. WSS was estimated using the Poiseuille parallel-plate approximation^27^, valid for high-aspect-ratio rectangular channels (w >>h). For the standard channel geometry (width w = 9.5 mm, height h = 365 µm), the width-to-height ratio (w/h ≈ 26) places the flow well within this limit, in which the velocity profile across the width is uniform except within ∼1 channel height of each side wall. Under this approximation, WSS relates directly to the measured centerline velocity as

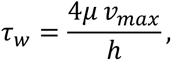

where *τ*_*w*_ is the wall shear stress, *μ* the dynamic viscosity of the perfusate, ℎ the channel height, and *v*_*max*_ the mid-height centerline velocity. Computing WSS from the directly measured *v*_*max*_ in each geometry ensures the estimate remains valid as channel dimensions, and hence the aspect ratio, are varied. Viscosity was taken as *μ* = 0.001 Pa·s for the perfusate at ∼20 °C.

### Seeding cells and initiating flow

Endothelial cells were cultured in vendor-recommended media and used at or below passage 8. For seeding, cells were detached with 0.05% trypsin at room temperature for 2–3 min until fully dissociated, collected, and centrifuged at 250 × g for 4 min. The supernatant was removed and cells were resuspended at 1.2 × 10⁶ cells/mL. Prior to seeding, fluidic channels were coated with collagen (50 µg/mL) for 1 h at 37 °C, rinsed once with PBS (Step 2 in Supplementary Fig. 9). PBS inside channels were removed by vacuum aspirator prior to seeding. The cell suspension (∼45 µL per channel) was introduced into each channel, and devices were held in the incubator (37 °C, 5% CO_2_) for 30 min to allow cell attachment (Step 3 in Supplementary Fig. 9). Each dish was then flooded with 10 mL of culture medium and returned to the incubator overnight (Step 4 in Supplementary Fig. 9). Flow was initiated the following day. To initiate flow, the impeller was first placed gently in the center of the channel array (Step 5, Supplementary Fig. 9). The top bracket was then positioned over the impeller, ensuring concentric alignment of the central shaft (Step 6, Supplementary Fig. 9). Handling was aided by inserting a 200 µL pipette tip into the vent hole, where the tolerance fit and friction held the bracket securely. Residual air was evacuated by positioning a vacuum aspirator over the vent hole (Step 7, Supplementary Fig. 9), and the vent hole was then sealed with adhesive tape (Scotch Transparent Tape, 3M, Minnesota, USA) to complete the assembly (Step 8, Supplementary Fig. 9).

### Immunofluorescence staining

The impeller and top bracket were removed from the dish, leaving only the channel array. Culture medium was aspirated, and the channels were rinsed once with PBS, rocking the dish several times to ensure the PBS flushed through the channels. Cells were then fixed by adding 5 mL of 4% paraformaldehyde (PFA) and rocking for 15 min at room temperature. Channels were cut open with a No. 11 scalpel to expose the cell-seeded surface (Supplementary Fig. 6B–C). Cells were permeabilized with 0.1% Triton X-100 for 10 min and blocked with 3% BSA for 1 h at room temperature. Primary antibodies against VE-cadherin (1:400; cat. no. 2500, Cell Signaling Technology, MA, USA) and GM130 (1:400; cat. no. 610823, BD Biosciences, CA, USA) were applied and incubated overnight at 4 °C. Samples were rinsed three times in PBS, then incubated for 1 h at room temperature with Alexa Fluor-conjugated secondary antibodies (1:500; cat. no. A-31573 and A-21422, Thermo Fisher, MA, USA) and Hoechst 33342 (1:1000; cat. no. H3570, Thermo Fisher, MA, USA). Samples were rinsed three times in PBS before imaging.

### Quantitative PCR

The impeller and top bracket were removed from the dish, leaving only the channel array. Culture medium was aspirated, and the channels were rinsed once with PBS, rocking the dish several times to ensure the PBS flushed through the channels. Channels were then cut open with a No. 11 scalpel to expose the cell-seeded surface (Supplementary Fig. 6B-D). RNA was extracted using the RNeasy Mini Kit (Qiagen, Hilden, Germany) according to the manufacturer’s protocol. Briefly, 350 µL of Buffer RLT (supplemented with 10 µL β-mercaptoethanol per 1 mL RLT) was pipetted onto the exposed cell-seeded surface and distributed across all eight channels of the array. Cells were gently scraped from the surface using the pipette tip, and the lysate was collected into a 1.5 mL Eppendorf tube. The lysate was disrupted and homogenized by passing it through a 21 G needle and syringe. An equal volume of 70% ethanol was added, and the resulting 700 µL sample was transferred to an RNeasy Mini spin column. Subsequent washing and elution steps followed the manufacturer’s protocol. Total RNA was extracted using the RNeasy Mini Kit and reverse-transcribed to cDNA using (qScript™ cDNA SuperMix, Part #95048-100, Quantabio, MA, USA). Quantitative PCR was performed on a QuantStudio 7 (Applied Biosystems, MA, USA) using SYBR Green master mix (Cat. No, #A25776, Applied Biosystem, MA, USA). Relative gene expression was calculated by the 2^(−ΔΔCt) method, normalized to PBGD and expressed relative to static controls, with primer specificity confirmed by melt-curve analysis. The following primers (5′→3′, forward and reverse) were used: KLF2 (CCTCACTTCCGTTGCTGATGA; GTCGAGGGGCCAACATTCTG), KLF4 (AGATGGGGTCTGTGACTGGA; CCTCCCCCAACTCACGGATA), NOS3 (TAACTCTGGCGCACTTGGG; AAGAAGTGGCATCAAGCGGA), THBD (CTGTGTGTCCGGTGTATGGT; ATGAGAGCCACAAACTGAGACC), EDN1 (GCTTTGTTTTGCCTGTCAAGG; TAATCACGTTGGCCCACTCC), SELE (GTGAAGCTCCCACTGAGTCC; AGCCAGAGGAGAAATGGTGC), VCAM1 (CTCCTGAGCTTCTCGTGCTC; TGACCCCTTCATGTTGGCTT), CCL2 (CAGCCAGATGCAATCAATGCC; TGGAATCCTGAACCCACTTCT), and PBGD (CATGTCTGGTAACGGCAATG; GTACGAGGCTTTCAATGTTG).

### Calcein AM viability staining

The impeller and top bracket were removed from the dish, leaving only the channel array. Culture medium was aspirated, and the channels were rinsed once with PBS, rocking the dish several times to ensure the PBS flushed through the channels. Channels were then cut open with a No. 11 scalpel to expose the cell-seeded surface (Supplementary Fig. 6B–D). Calcein AM (2 µM in culture medium) was added onto the cell surface and incubated for 10 min. The channels were rinsed once with PBS, then covered with fresh medium and imaged immediately (Echo Revolve microscope, BICO, Gothenburg, Sweden).

### Cell segmentation and alignment analysis

Fluorescence microscopy images were acquired as multi-channel z-stack CZI files using a confocal microscope (LSM 980, Zeiss, Oberkochen, Germany). Each CZI file contained four channels: phalloidin-labelled F-actin (488 nm), GM130 (568 nm), VE-Cadherin (647 nm), and DAPI (405 nm). Prior to segmentation, images were pre-processed using a custom Python pipeline. Briefly, maximum intensity z-projections were generated independently for the VE-Cadherin and DAPI channels using the aicspylibczi library (v0.x). Each projected channel was subjected to adaptive histogram equalisation (scikit-image equalize_adapthist, clip_limit = 0.02– 0.05) with channel-specific parameters optimised per cell type to account for differences in staining intensity across endothelial cell subtypes. The processed channels were exported as two-channel RGB TIFF files with VE-Cadherin assigned to the red channel and DAPI assigned to the green channel. Cell segmentation was performed using Cellpose 3 (v3.1.1) with the cyto3 model as the initialisation point for fine-tuning. To generate training data, a representative subset of images spanning all six endothelial cell subtypes (BMVEC, DMVEC, HAEC, HCAEC, HSaVEC, HUVEC) and all shear stress conditions (static, 2, 15, and 40 dyn/cm²) were manually annotated using the Cellpose graphical user interface. Cell boundaries were traced individually for a total of approximately 500 cells across four images, with particular attention paid to correctly delineating elongated cells at high shear stress and cells with faint junction staining.

The cyto3 model was fine-tuned on these annotations using stochastic gradient descent (learning rate = 0.005, weight decay = 0.0001, 100 epochs), with VE-Cadherin designated as the cytoplasm channel and DAPI as the nuclear channel. The resulting custom model was applied to all images using a cell diameter of 80 pixels for cobblestone morphology cell types (BMVEC, HAEC, HCAEC, HSaVEC, HUVEC) and 35 pixels for the highly elongated DMVEC, with a flow threshold of 0.8 and cell probability threshold of −2.0 to maximize detection sensitivity. Segmentation outputs were quality-controlled by visual inspection of all images. Each detected region was required to contain exactly one DAPI nucleus centroid, verified programmatically by overlaying the Cellpose masks with a DAPI nucleus segmentation (also performed using Cellpose with the nuclei model, diameter = 34 pixels). Regions containing zero or more than one nucleus were excluded, rejecting false positives from background noise and merged cell detections respectively. Cell orientation angles were measured relative to the vertical axis of the image, where 0° corresponds to a vertically oriented cell and 90° corresponds to a horizontally oriented cell aligned with the direction of flow. Semi-rose plots display orientation data as axial distributions in a full 180° circular format. The plot is oriented with 0° at the top (vertical) and flow direction (90°).

### Golgi Orientation Analysis

Golgi polarization relative to the nucleus was quantified from confocal immunofluorescence images using a custom Python-based image analysis pipeline. Z-stack images acquired in the DAPI and GM130 channels were maximum-intensity projected along the z-axis prior to analysis. Nuclear segmentation was performed using Cellpose (v2.0) with the built-in “nuclei” model applied to the DAPI channel, with cell diameter estimated automatically and flow and cell probability thresholds set to 0.4 and 0.0, respectively. Cell body boundaries were defined from the VE-cadherin channel segmentation mask described above, and used to spatially assign each nucleus and Golgi structure to an individual cell. Golgi structures were segmented from the GM130 channel by applying Otsu automatic thresholding to a Gaussian-smoothed image (σ = 1.5 px), followed by morphological filtering to remove small debris (minimum area = 120 px²). To establish a strict one-to-one correspondence between each nucleus and its associated Golgi apparatus, a Golgi-first nearest-neighbor assignment was employed: each Golgi blob was first assigned to the nearest nucleus centroid within the same cell boundary, and where multiple Golgi blobs were assigned to a single nucleus, only the largest was retained. This strategy excluded small peri-nuclear GM130 puncta from the directional measurement. Golgi orientation was defined as the angle of the vector from the nucleus centroid to the Golgi centroid, measured clockwise from the axis perpendicular to flow (0° = perpendicular to flow; 90° = downstream). Per-cell angles were aggregated across a minimum of two biological replicates per condition and analyzed using circular statistics, including the mean resultant vector angle and the Rayleigh R statistic as a measure of directional coherence (R = 0, uniformly random; R = 1, perfectly aligned). Angle distributions were visualized as rose plots.

### Isolating lysate for bulk RNA-sequencing

The impeller and top bracket were removed from the dish, leaving only the channel array. Culture medium was aspirated, and the channels were rinsed once with PBS, rocking the dish several times to ensure the PBS flushed through the channels. Channels were then cut open with a No. 11 scalpel to expose the cell-seeded surface (Supplementary Fig. 5B-D). DNA/RNA Shield (250 µL; #R1100, Zymo Research, Irvine, CA, USA) was added onto the exposed cell-seeded surface, and cells were scraped using the tip of a 200 µL pipette. The collected lysate was shipped to Plasmidsaurus (South San Francisco, CA, USA) for bulk RNA sequencing.

### Principal Component Analysis

Bulk RNA-sequencing count data were processed in R. Raw count matrices were extracted from a tab-delimited expression file and sample identifiers were standardized using a predefined mapping table to ensure consistent labeling across analyses. Gene-level counts were filtered to remove lowly expressed genes, retaining genes with counts per million (CPM) ≥1 in at least one sample. Library size normalization was performed using the trimmed mean of M-values (TMM) method implemented in the edgeR framework. Normalized expression values were transformed to log2-CPM with a prior count of 1. Principal component analysis (PCA) was performed on the filtered log2-CPM matrix using the prcomp function in R, with centering and scaling applied across genes. The input matrix was transposed such that rows corresponded to samples and columns to genes. The first two principal components (PC1 and PC2) were extracted and used for visualization. For visualization, samples were annotated by cell type and experimental condition based on standardized sample names. PCA plots were generated using ggplot2, with points colored by cell type and shaped by experimental condition. Elliptical contours representing the 95% confidence region of each condition group were computed based on the covariance structure of PC1 and PC2 and overlaid to aid interpretation of group-level clustering.

### Differential gene expression analysis

Differential expression analysis was performed using the edgeR and limma pipelines. To account for baseline transcriptional differences between endothelial cell lines, differential expression analyses were performed using a linear modeling framework that included both cell-line and condition effects. The design matrix was constructed using the formula:∼cellline_factor+condition_factor, where the tested coefficient represented the flow condition effect adjusted for baseline cell-line differences. Genes with insufficient expression were filtered using edgeR::filterByExpr based on the experimental design matrix. Library size normalization was performed using the trimmed mean of M-values (TMM) normalization method implemented in edgeR. Normalized count data were transformed using the limma voom procedure to estimate the mean–variance relationship and generate precision weights for linear modeling. Linear models were fitted using limma::lmFit followed by empirical Bayes variance moderation using limma::eBayes. Differential expression statistics were extracted using limma::topTable. For each gene, the moderated t-statistic was used as the signed ranking metric for downstream gene set enrichment analysis (GSEA). Additional output metrics included log2 fold change (logFC), average expression (AveExpr), raw P values, adjusted P values using the Benjamini–Hochberg false discovery rate (FDR) correction, and B-statistics. Volcano plots were generated using ggplot2 and ggrepel. Genes were classified as upregulated or downregulated using thresholds of FDR < 0.05 and absolute log2 fold change ≥1. The top-ranked genes by adjusted P value were labeled on the volcano plots. Expression heatmaps were generated using the pheatmap package. For each differential expression comparison, the top 30 genes ranked by adjusted P value were selected from the limma-voom differential expression results. Expression values were standardized by row using z-score transformation to facilitate visualization of relative expression patterns across samples. Samples were annotated according to both flow condition and endothelial cell line. Hierarchical clustering was applied to both rows and columns using default pheatmap clustering settings.

### Validation with publicly available datasets

For both GSE151867 and GSE294621 datasets, gene-level counts were filtered using edgeR::filterByExpr, normalized using the trimmed mean of M-values (TMM) method, and modeled using the same limma-voom framework. Differential expression thresholds (FDR < 0.05, |log2 fold change| > 1) were applied to the platform’s own dataset, to ensure comparability across all three. The static versus 15 dyn/cm² contrast was used for comparison with GSE151867, and the static versus 10 dyn/cm² contrast was used for comparison with GSE294621. Concordance between the platform’s HUVEC dataset and each public dataset was assessed in three complementary ways. First, the percentage of the platform’s differentially expressed genes (DEGs) that were also identified as DEGs in each public dataset, considering DEGs in either direction, was calculated. Second, Spearman correlation of log2 fold-change values between the platform’s dataset and each public dataset was calculated across two gene sets: all genes shared between the two ranked gene lists and genes meeting the DEG threshold in both datasets. Third, a canonical panel of laminar shear-responsive genes (*KLF2*, *KLF4*, *NOS3*, *THBD*, *VCAM1*, *ICAM1*, *SELE*, *CCL2*, *EDN1*, *HMOX1*, *CAV1*, *FN1*, and *SERPINE1*) was examined directly across the platform’s dataset and each public dataset, reporting log2 fold-change, adjusted P value, and direction of regulation for each gene, with concordance defined as agreement in direction of regulation between the platform’s dataset and the corresponding public dataset.

### Gene Set enrichment analysis

Gene set enrichment analysis was performed using the fgsea package with the fgseaMultilevel algorithm. Ranked gene lists were generated from the moderated t-statistics obtained from the cell-line-adjusted limma-voom models. Gene Ontology Biological Process (GO:BP) pathways were obtained from the Molecular Signatures Database (MSigDB) using the msigdbr package. For each comparison, enrichment analysis was performed using a minimum pathway size of 10 genes and a maximum pathway size of 500 genes. Analyses were run using a fixed random seed and single-threaded execution to improve reproducibility. Enrichment results included normalized enrichment scores (NES), adjusted P values, pathway sizes, and leading-edge gene subsets. For visualization, the top 10 positively enriched pathways and top 10 negatively enriched pathways ranked by adjusted P value were selected for each comparison. Dot plots were generated using ggplot2, with normalized enrichment score represented on the x-axis, pathway names on the y-axis, point size corresponding to pathway size, and point color corresponding to NES direction and magnitude.

### Graded response analysis

To determine whether shear-responsive genes followed a discrete, condition-specific pattern or a continuous, dose-dependent trajectory, mean log2(CPM+1) expression was calculated for each gene at each of the four shear conditions (static, 2, 15, and 40 dyn/cm²), averaging across all replicates and endothelial subtypes within each condition. For each gene, Spearman correlation was calculated between mean expression and ranked shear magnitude (static = 1, 2 dyn/cm² = 2, 15 dyn/cm² = 3, 40 dyn/cm² = 4). Genes with an absolute Spearman correlation coefficient (|ρ|) ≥ 0.9 were classified as shear-magnitude-dependent, with the sign of the correlation determining classification as shear-induced or shear-repressed. To confirm that these graded trends reflected genuine differential expression rather than correlation alone, shear-magnitude-dependent genes were additionally required to reach statistical significance (FDR < 0.05) in the cell-line-adjusted limma-voom pairwise contrast corresponding to their trajectory shape. Genes meeting the spearman threshold were first classified by trajectory shape: those whose expression changed by less than a log2 fold-change of 0.5 between 15 and 40 dyn/cm² were classified as plateau genes, reaching their maximal response by 15 dyn/cm², genes whose expression changed by a log2 fold-change of 0.5 or more between these two conditions were classified as progressive genes, whose expression continued to change between 15 and 40 dyn/cm². Progressive genes were required to reach significance in both the 40 dyn/cm² versus static contrast and the 40 dyn/cm² versus 15 dyn/cm² contrast. Genes satisfying both the Spearman threshold and the corresponding pairwise significance criterion were classified as high-confidence shear-magnitude-dependent genes, comprising shear-induced and shear-repressed subsets. To visualize these trajectories, expression values for each gene were z-score normalized across all samples and averaged within each condition to generate group-level mean trajectories with associated standard error, stratified by direction of response (induced or repressed). Individual gene-level trajectories were overlaid on group-level trajectories to illustrate the consistency of directional response across the full shear range.

### Biological contrast

Biological contrast analyses were performed using predefined endothelial response groups derived from flow-alignment behavior. Endothelial cell lines were categorized into low-shear aligners (DMVEC and HsaVEC) and low-shear non-aligners (HUVEC, HCAEC, HAEC, and BMVEC). Pairwise and interaction-style contrasts were designed to capture biologically meaningful transcriptional responses associated with endothelial mechanoadaptation. Biological contrasts included: (i) low-shear aligners at 2 dyn/cm² relative to static controls, (ii) low-shear non-aligners at 2 dyn/cm² relative to static controls, (iii) an interaction-style low-shear alignment contrast comparing the 2D response between aligners and non-aligners, (iv) a threshold transition contrast comparing 15 dyn/cm² versus 2 dyn/cm² within non-aligning endothelial populations, and (v) a reinforcement contrast comparing 40 dyn/cm² versus 15 dyn/cm² across all endothelial cell lines. Differential expression analysis and gene set enrichment analysis was performed as described above. To reduce redundancy and facilitate systems-level interpretation of enriched GO Biological Process pathways, significant pathways were collapsed into higher-order biological signature groups using deterministic keyword-based classification frameworks.

### Software and data analysis

All cell segmentation analysis was performed in Python 3.10 using NumPy, scikit-image, SciPy, and Matplotlib. All golgi orientation analyses were performed in Python (Cellpose, scikit-image, SciPy, NumPy, Matplotlib). All RNA-sequencing analyses were performed in R using packages including edgeR, limma, fgsea, msigdbr, ggplot2, pheatmap, ggforce, cowplot, ggrepel, dplyr, tidyr, tibble, stringr, and readr.

## Data Availability

RNA-sequencing data is deposited in the Gene Expression Omnibus under accession GSE339144. During the review process, data can be accessed via the following review token: uxyxeuiajfkdtmv. Data will be made publicly available upon publication. Source data underlying all figures are provided as supplementary files. CAD and cutting files, a bill of materials, and an assembly protocol for the PROPEL impeller and channel array are included in the Zenodo deposit (DOI: 10.5281/zenodo.21420629).

## Code Availability

Analysis scripts, hardware design files, bills of materials, assembly protocols, and custom-trained Cellpose models for VE-cadherin/DAPI cell-body segmentation are available on Zenodo (DOI: 10.5281/zenodo.21420629) and are actively maintained at https://github.com/PROPELflow/PROPELflow-platform.

## Acknowledgements

We thank Dr. Ksenia Safina and Prof. Peter van Galen for their valuable input on bioinformatic analysis strategies, colleagues who provided generous gifts of reagents, and our lab members for their support and good counsel.

