## Supplementary Figures for "PROPEL: a high-throughput shear-stress platform reveals organotypic thresholds in endothelial mechano-adaptation"

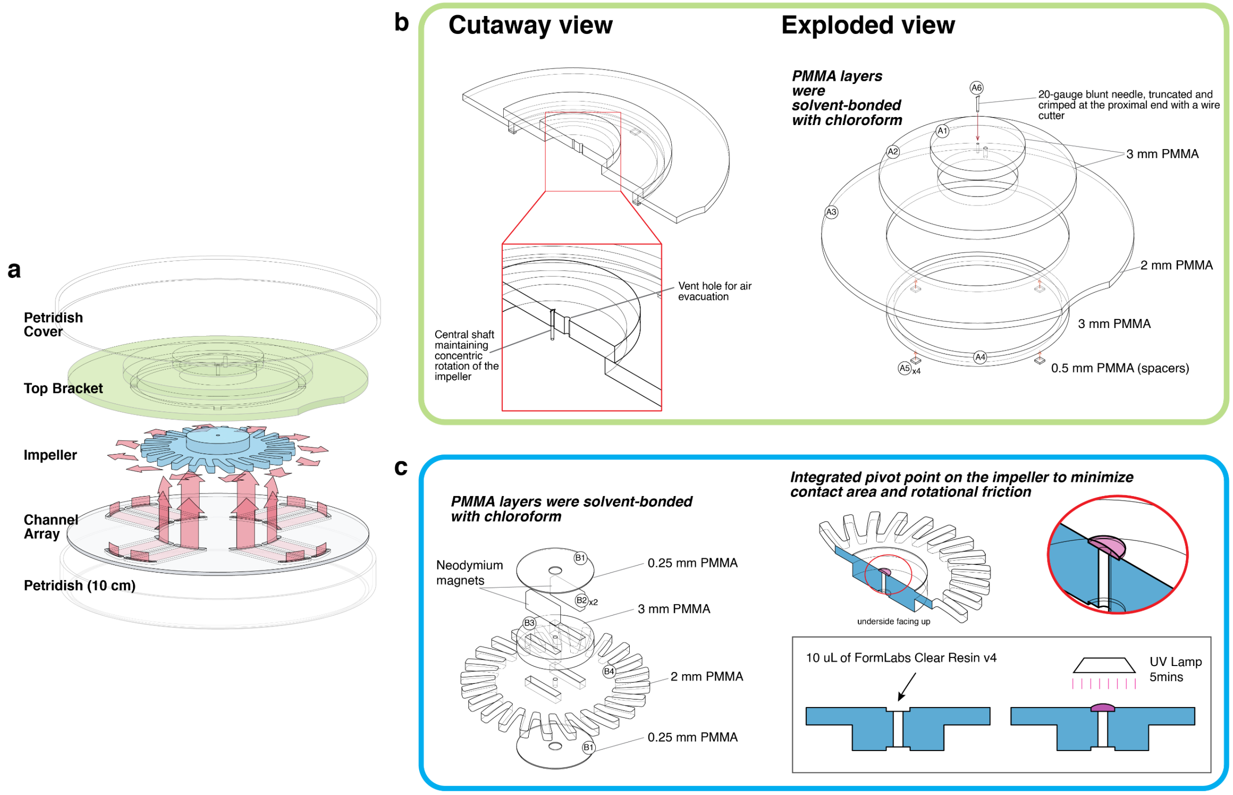


Supplementary.Figure.7¡.Fabrication.of.the.top.bracket.and.impeller¡ (a) *Exploded view of the components comprising PROPEL*. (b) *Exploded and cutaway views of the top bracket, fabricated entirely from PMMA sheets of varying thickness (0.25, 2, and 3 mm) solvent-bonded with chloroform*. (c) *Exploded and cutaway views of the impeller, assembled from solvent-bonded PMMA sheets that seal a pair of permanent magnets inside. A pivot point was added by UV-curing a droplet of resin on the underside, minimizing friction during rotation.*

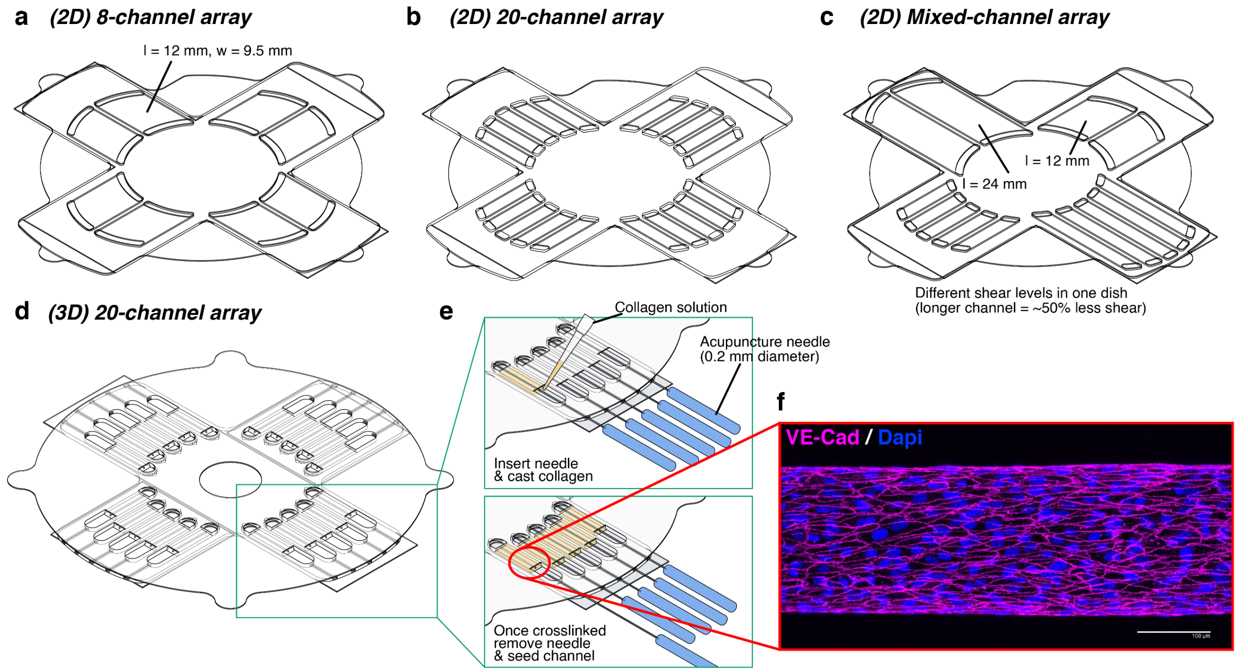


Supplementary.Figure.8¡.Channel.arrays.of.varying.configuration?dimension?.and.geometry¡ (a) *An 8-channel array (each channel l = 12 mm, w = 9.5 mm)*. (b) *A 20-channel array (each channel l = 12 mm, w = 3 mm)*. (c) *A mixed-dimension array combining channels of four geometries in a single dish (2× l = 12 mm/w = 9.5 mm; 2× l = 24 mm/w = 9.5 mm; 5× l = 12 mm/w = 3 mm; 5× l = 24 mm/w = 3 mm); because shear scales with channel dimension, each channel type experiences a distinct shear stress, enabling multiple shear levels within one dish*. (d) *An array of 20 three-dimensional cylindrical channels*. (e) *Fabrication of cylindrical channels using an acupuncture needle as a sacrificial mold: needles are inserted into the channel array, uncrosslinked collagen is pipetted in, and after crosslinking the needle is withdrawn to leave a hollow lumen for cell seeding*. (f) *Representative micrograph of a cylindrical channel seeded with HUVECs, stained for VE-cadherin and DAPI*.

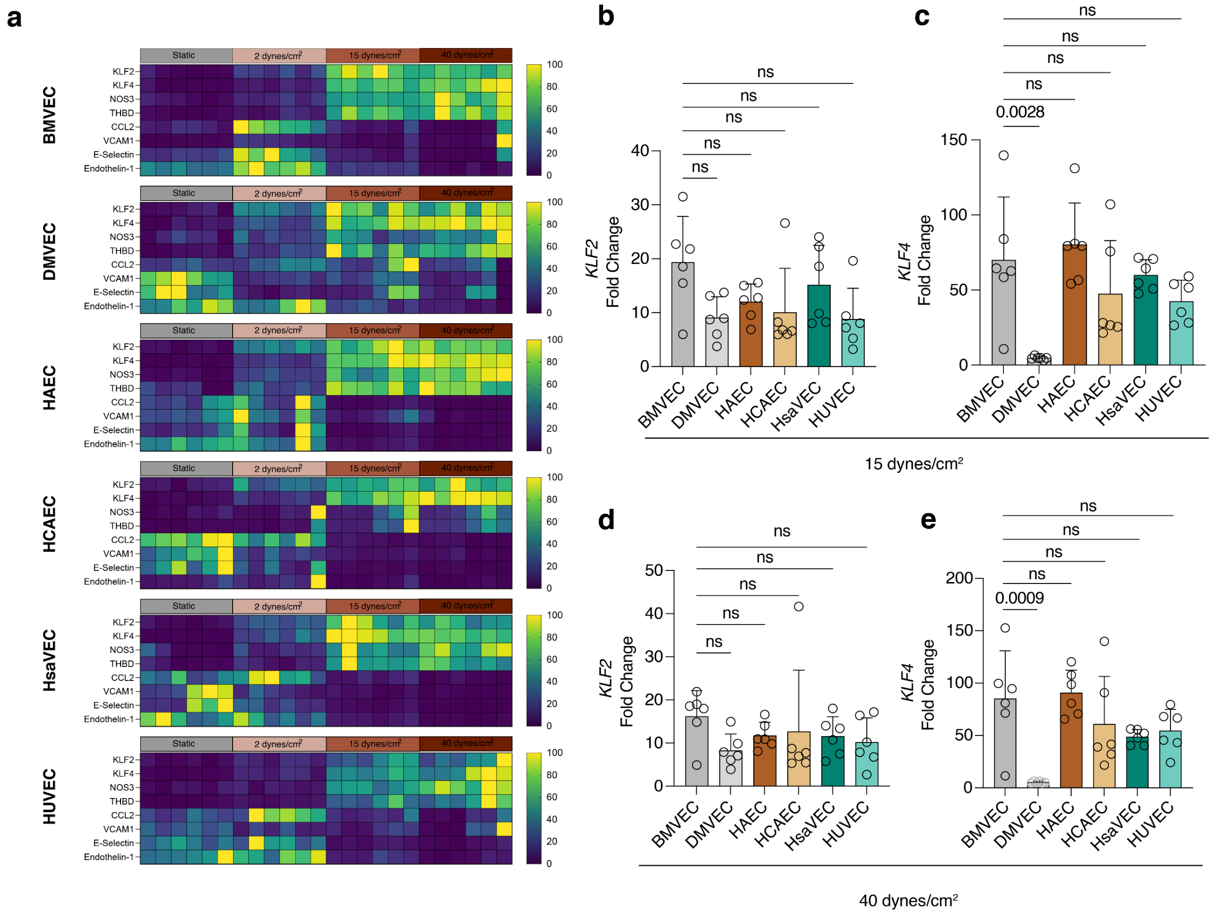


Supplementary.Figure.9¡.(a) *Heatmaps of normalized relative expression for KLF2, KLF4, NOS3, THBD, CCL2, VCAM1, SELE, and ET-1 across six endothelial subtypes (BMVEC, DMVEC, HAEC, HCAEC, HSaVEC, HUVEC) under static, 2, 15, and 40 dyn/cm² conditions. Each column represents an independent biological replicate.* (b- e) *Relative gene expression of KLF2* (b?d) *and KLF4.*(c?e) *across the six endothelial subtypes subjected to 15 dyn/cm²* (b?c) *or 40 dyn/cm²* (d?e) *for 3 days (N = 6 per subtype).*

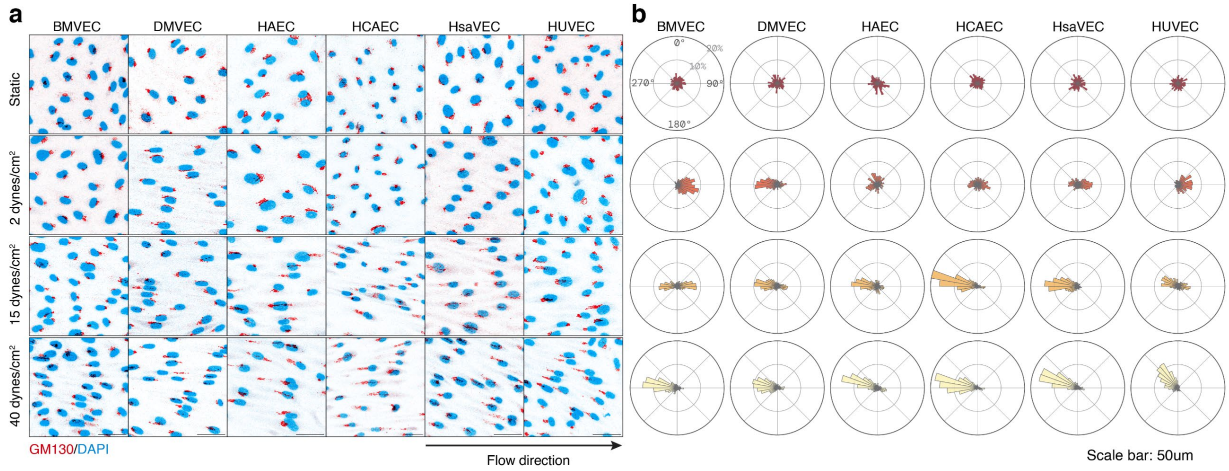


Supplementary.Figure.0¡.(a) *Representative immunofluorescence images of GM130 (red) and DAPI (blue) in each of the six endothelial subtypes under static, 2 dyn/cm$^2$, 15 dyn/cm$^2$, and 40 dyn/cm$^2$ conditions for 3 days*. (b) *Polar histograms of golgi-nucleus polarization angle relative to flow direction for each subtype under static, 2 dyn/cm$^2$, 15 dyn/cm$^2$, and 40 dyn/cm$^2$ conditions for 3 days*.

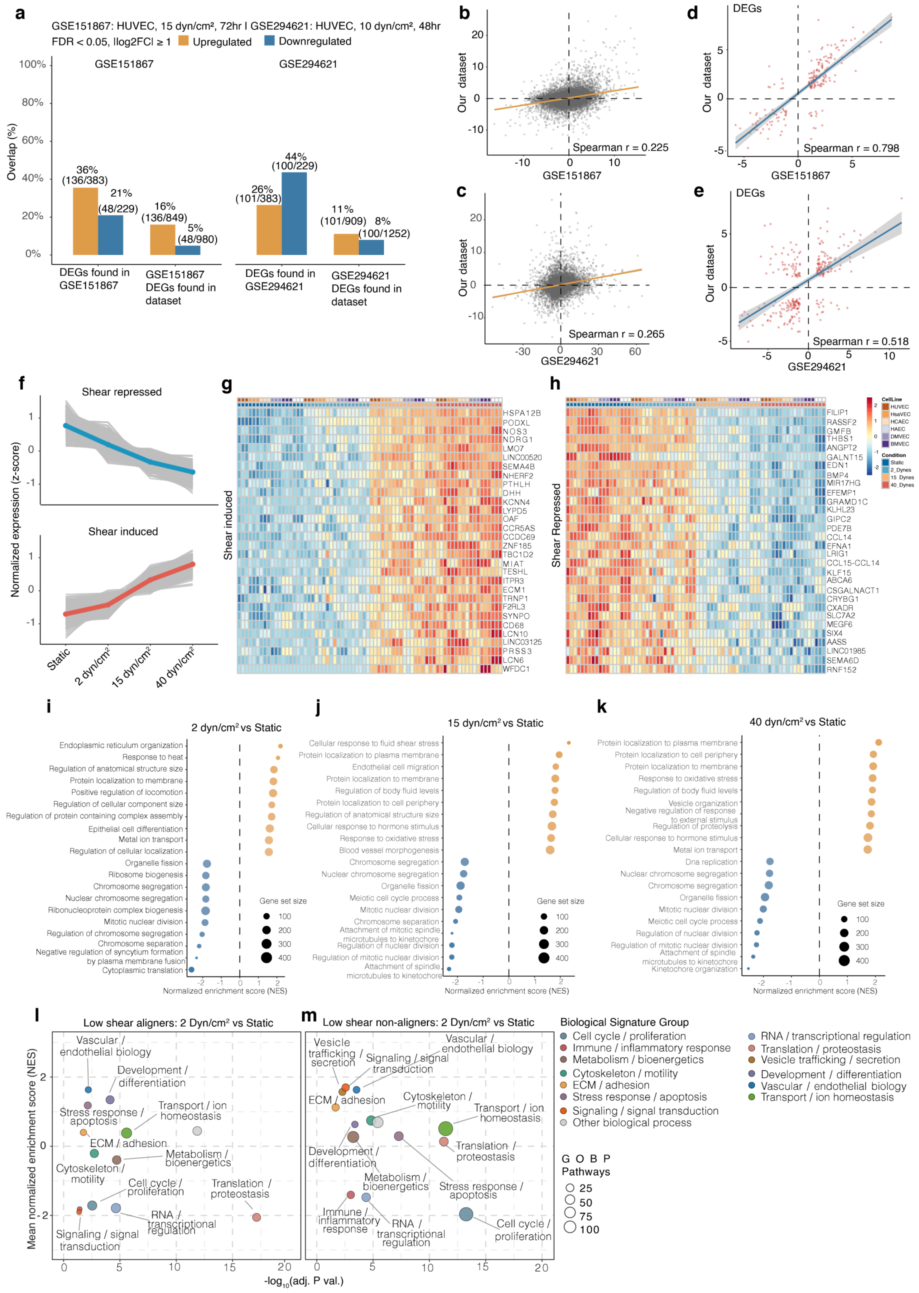


Supplementary.Figure.❶ Supplementary.Figure.X¡.Validation.of.RNA_seq.findings.against.public.datasets.and. graded.transcriptional.response.to.shear.magnitude¡ (a) *Percentage overlap of differentially expressed genes (DEGs) between this dataset and two publicly available shear stress datasets, GSE151867 (15 dyn/cm², 72 h) and GSE294621 (10 dyn/cm², 48 h).* (b?c) *Correlation of $\log_2$ fold-change values across all genes shared between this dataset and GSE151867* (b) *or GSE294621* (c). (d?e) *Correlation of $\log_2$ fold-change values restricted to genes differentially expressed in both datasets, for this dataset versus GSE151867* (d) *or GSE294621* (e). (f) *Normalized expression trajectories of shear-repressed (top) and shear-induced (bottom) genes across static, 2, 15, and 40 dyn/cm² conditions. Colored lines indicate group means; grey lines indicate individual genes.* (g?h) *Heatmaps of normalized expression for representative shear-induced* (g) *and shear-repressed* (h) *genes.* (i-k) *Top ten enriched (orange) and top ten repressed (blue) Gene Ontology Biological Process (GO BP) pathways for the 2* (i)*, 15* (j)*, and 40 dyn/cm²* (k) *versus static comparisons, ranked by normalized enrichment score. Bubble size indicates gene set size.* (l?m) *GO BP pathways grouped into biological signature categories for the 2 dyn/cm² versus static comparison, in low-shear aligner* (l) *and low-shear non-aligner* (m) *endothelial subtypes. Bubble size indicates the number of GO BP pathways contributing to each signature group.*

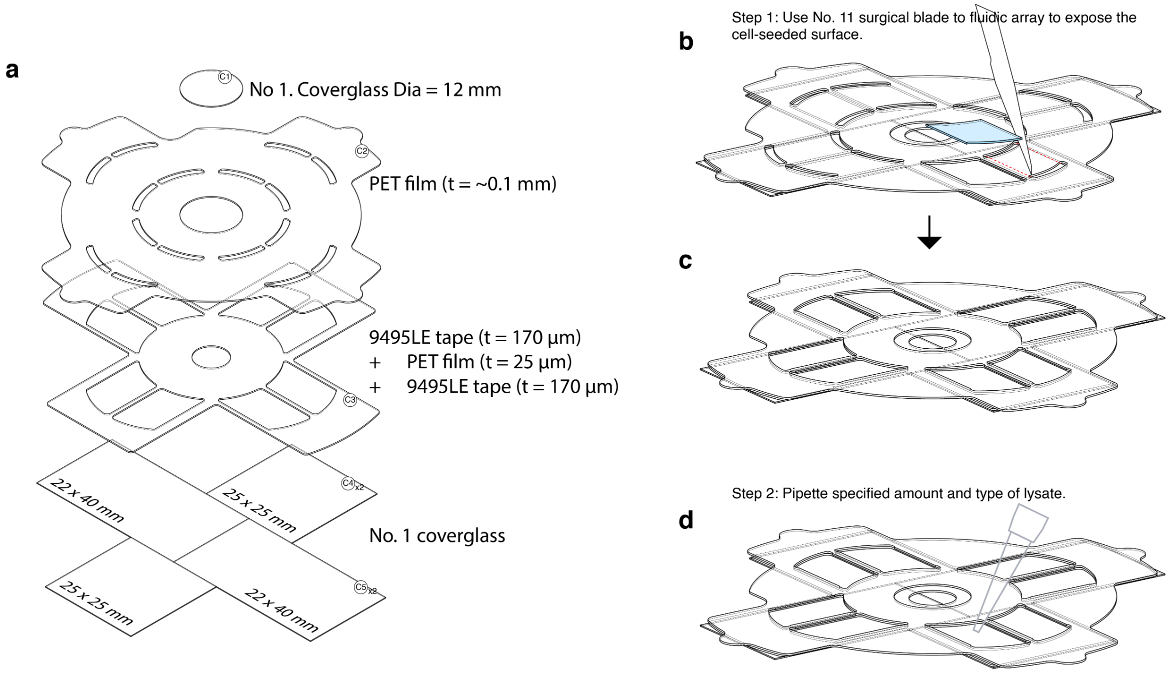


Supplementary.Figure.❷¡.Fluidic.channel.array.assembly.and.cell.lysis.workflow¡ (a) *Exploded view of the layers comprising the fluidic channel array.* (b- c) *Schematic of the workflow for lysing cells within the channel array for qPCR and bulk RNA sequencing.* (b- c) *A No. 11 surgical blade is used to cut open the top flap of the channel array, exposing the cell-seeded surface.* (d) *Lysis buffer (e.g., Buffer RLT or DNA/RNA Shield) is pipetted directly onto the exposed cells for collection.*

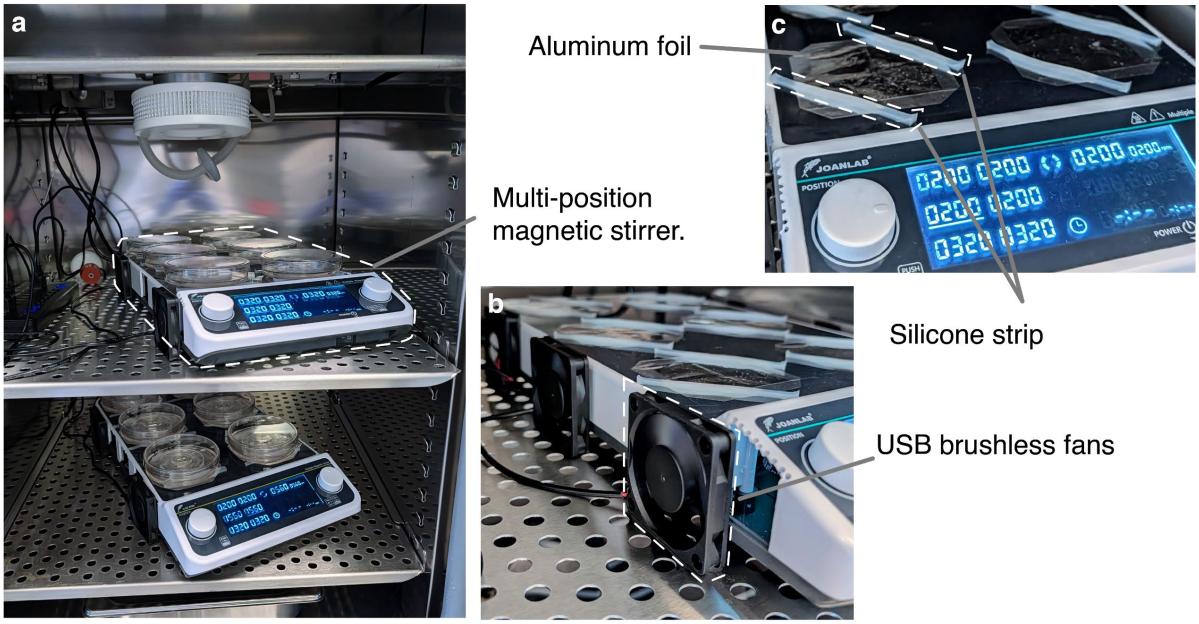


Supplementary Figure 3. Magnetic stirrer modifications for long-term operation. (a) *Multi-position magnetic stirrer blocks operating inside the incubator, with PROPEL-integrated petri dishes mounted on each stirrer position*. (b) *USB computer fans attached to the side of a stirrer block to improve cooling of the internal electronics*. (c) *Close-up showing the placement of the aluminum foil (radiative heat shield) and silicone strips (dish standoff and friction grip)*.

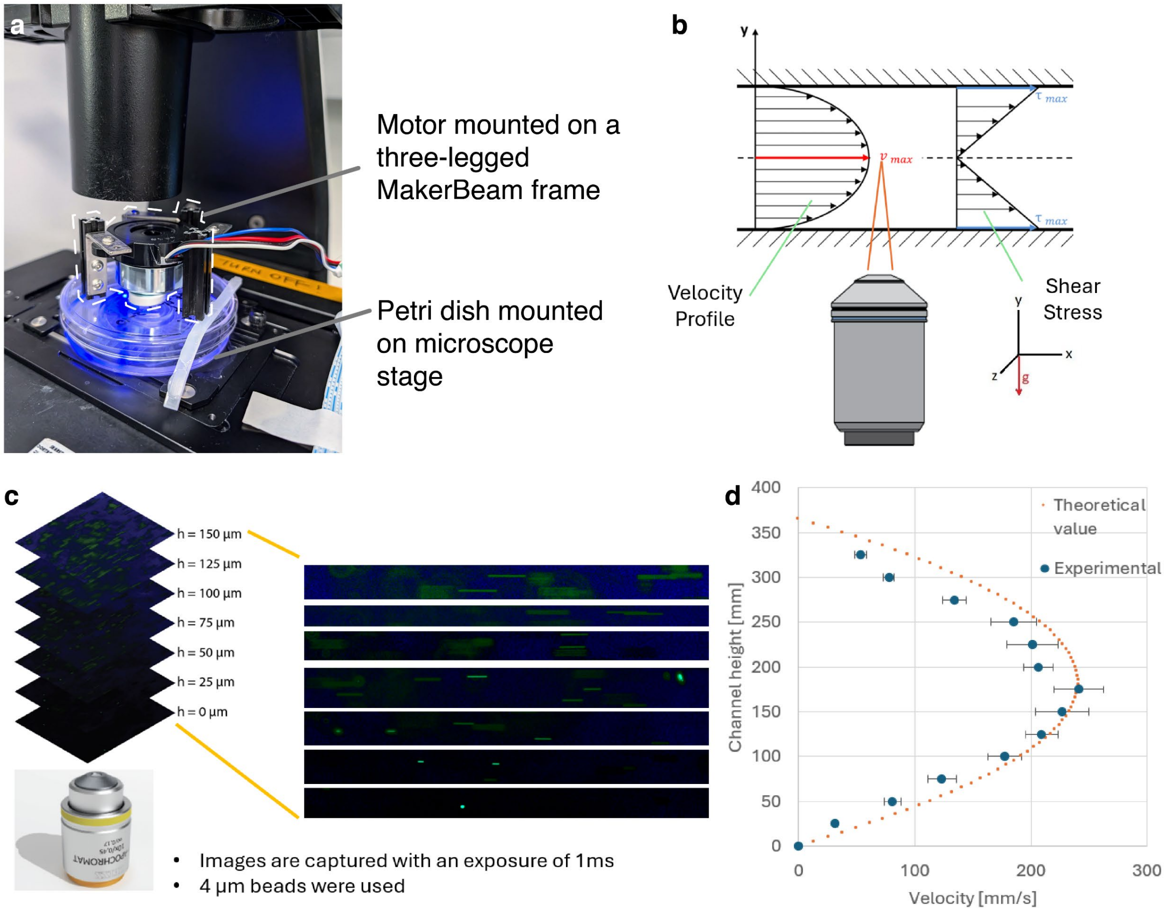


Supplementary.Figure.❹.Flow.characterization.by.particle.streak.velocimetry¡ (a) *Photograph of the top-mounted magnetic stirrer, built from a three-legged MakerBeam frame, resting on the lid of a PROPEL-integrated petri dish to drive the impeller from above during imaging.* (b) *Schematic of the imaging plane within the channel, together with the expected velocity profile and wall shear stress distribution for parallel-plate flow.* (c) *Particle streak velocimetry: a 1 ms exposure captures each bead as a streak whose length equals the distance the bead travels in 1 ms, providing a direct measure of local velocity.* (d) *Measured velocity profile across the channel (h = 365 µm, w = 9.5 mm) driven by PROPEL; the orange trace shows the theoretical parabolic profile for comparison.*

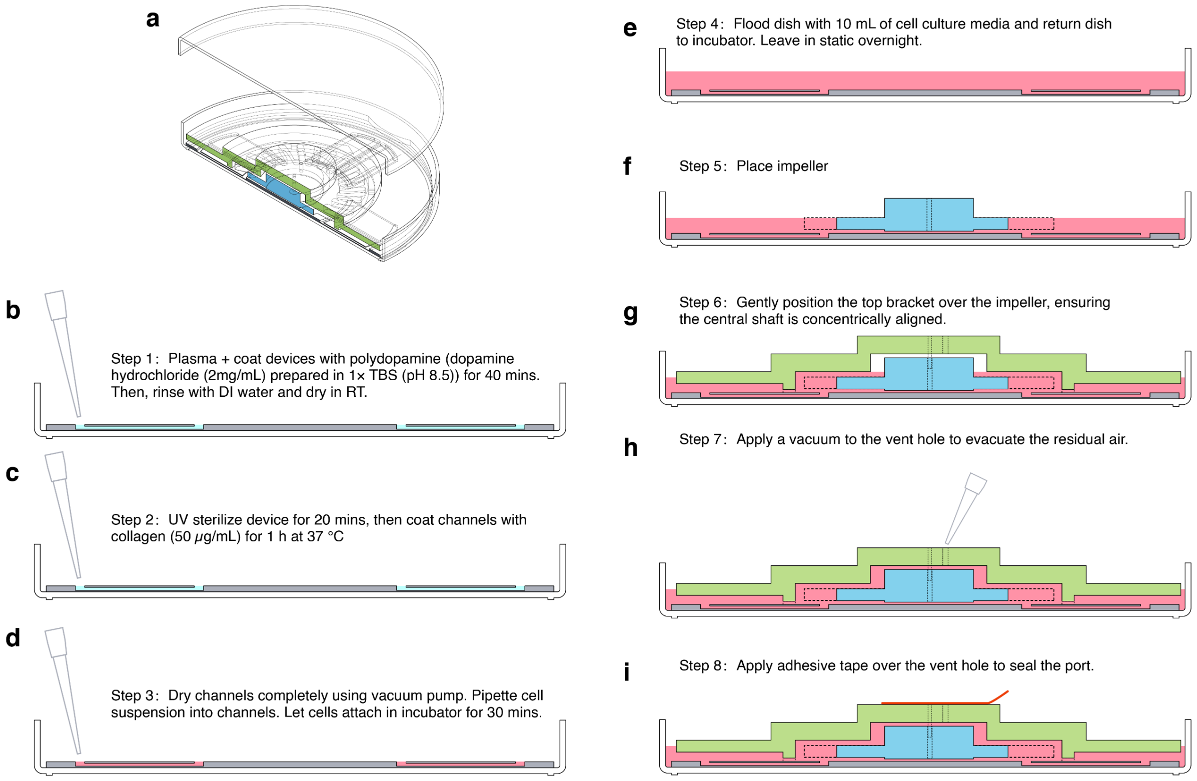


Supplementary.Figure.5¦ Device.coating?cell.seeding?and.flow.initiation¡.Schematic.of.the.full.workflow.from. surface.functionalization.through.flow.initiation¡ (a) *Cutaway view of the assembled PROPEL device*. (b) *Step 1: plasma treatment and polydopamine coating of the channel surface*. (c) *Step 2: collagen coating of the channel surface*. (d) *Step 3: drying the channels and seeding the cell suspension*. (e) *Step 4: flooding the dish with 10 mL of culture medium*. (f) *Step 5: placing the impeller onto the channel array*. (g) *Step 6: placing the top bracket over the impeller*. (h) *Step 7: evacuating residual air from the assembly*. (i) *Step 8: sealing the vent port with adhesive tape*.
